# Perfusion Quantification in the Human Brain Using DSC MRI – Simulations and Validations at 3T

**DOI:** 10.1101/2022.04.27.489686

**Authors:** J. Schulman, E.S. Sayin, A. Manalac, J. Poublanc, O. Sobczyk, J. Duffin, J.A. Fisher, D.J. Mikulis, K. Uludağ

**Affiliations:** Department of Medical Biophysics, University of Toronto, Toronto, ON, Canada; Techna Institute, University Health Network, Toronto, ON, Canada; Center for Neuroscience Imaging Research, Institute for Basic Science & Department of Biomedical Engineering, Sungkyunkwan University, Suwon, Republic of Korea; Department of Physiology, University of Toronto, Toronto, ON, Canada; Joint Department of Medical Imaging and the Functional Neuroimaging Lab, University Health Network, Toronto, ON, Canada; Department of Anaesthesia and Pain Management, University Health Network, University of Toronto, Toronto, ON, Canada; The Joint Department of Medical Imaging, The Toronto Western Hospital, Toronto, ON, Canada; Toronto General Hospital Research Institute, Toronto General Hospital, Toronto, ON, Canada

**Keywords:** Perfusion, DSC, gadolinium, deoxyhemoglobin, simulations, susceptibility

## Abstract

Gadolinium (Gd) and deoxyhemoglobin (dOHb) are paramagnetic contrast agents capable of inducing changes in T_2_*-weighted MRI signal, utilized in dynamic susceptibility contrast (DSC) MRI. With multiple contrast agents and analysis choices, there are a variety of questions as to its capability to accurately quantify perfusion values. To address these questions, we developed a novel signal model for DSC MRI that incorporates signal contributions from intravascular and extravascular water proton spins at 3T for arterial, venous, and cerebral tissue voxels. This framework allowed us to model the MRI signal in response to changes in Gd and dOHb concentrations, and the effects that various experimental and tissue parameters have on perfusion quantification. We compared the predictions of the numerical simulations with those obtained from experimental data at 3T on six healthy human subjects using Gd and dOHb boluses as contrast agents. Using standard DSC analysis, we identified perfusion quantification dependencies in the experimental results that were in close agreement with the simulations. We found that a reduced baseline oxygen saturation (base-S_a_O_2_), greater susceptibility of applied contrast agent (Gd *vs* dOHb), and larger magnitude of the hypoxic drop (ΔS_a_O_2_) reduces overestimation of the cerebral blood volume (*rCBV*) and flow (*rCBF*). Furthermore, shortening the bolus duration increases the accuracy and reduces the calculated values of mean transit time (*MTT*). This study demonstrates that changes in Gd and dOHb can be described by the same unifying theoretical framework, as validated by the experimental results. Based on our work, we suggest practices in DSC MRI that increase accuracy and reduce inter- and intra-subject variability. In uncovering a wide array of quantification dependencies, we argue that caution must be exercised when comparing perfusion values obtained from a standard DSC MRI analysis when employing different experimental paradigms.

## Introduction

Cerebral perfusion imaging, and the perfusion metrics quantified therein, plays an important role in investigating various disease states characterized by vascular abnormality, such as atherosclerotic disease, vasculitis, Moyamoya disease, and neoplasms (Copen *et al*., 2007; Harris *et al*., 1998; Law *et al*., 2008; Qiao *et al*., 2017). One such form of perfusion imaging is dynamic susceptibility contrast (DSC), wherein a paramagnetic contrast agent, typically gadolinium (Gd), is injected intravenously as a bolus to enhance tissue contrast in vascularized regions and enable perfusion quantification using MRI (Jahng *et al*., 2014).

With seven unpaired electrons, Gd is a powerful MRI contrast agent providing a strong signal change in T_2_*-weighted MR images throughout the brain (Jahng *et al*., 2014). The paramagnetic contrast agent creates inhomogeneities in the local magnetic field, diminishing the phase coherence and increasing the relaxation rate 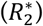 of extravascular and intravascular water proton spins – this decreases the overall MR signal within the region (Jahng *et al*., 2014). The observed signal and relaxation rate time courses resulting from susceptibility contrast agent can then be used to calculate the signal peak, time to peak, cerebral blood volume (*CBV* (ml/100g)), cerebral blood flow (*CBF* (ml/100g/min)), and mean transit time (*MTT* (s)), all of which have clinical and research utility (Copen *et al*., 2007; Harris *et al*., 1998; Law *et al*., 2008; Qiao *et al*., 2017). Relative *CBF* and *CBV (rCBF* and *rCBV*) for each tissue voxel is typically quantified in reference to the arterial input function (AIF), as measured over an arterial voxel, to reduce inter-subject variability that arises from varying bolus duration, rate of injection, and concentration (Calamante, 2013).

While Gd is robust in generating a quantifiable signal change, its use has been associated with notable limitations regarding safety. Gd is a toxic metal and must be chelated with an organic compound that prevents interaction with the tissue (Schlaudecker and Bernheisel, 2009). While newer macrocyclic Gd-chelates are safer than linear Gd-chelates (Matthias *et al*., 2011; Uhlig *et al*., 2020), they have still been shown to cause minor side effects following administration. As well, Gd is naturally invasive as it must be injected intravenously. While major complications are rare, they are more common in patients with renal disease; about 4% of those with renal disease who receive Gd contrast develop nephrogenic systemic fibrosis, which has a mortality rate of 31% (Schlaudecker and Bernheisel, 2009; Voth *et al*., 2011). There is also growing evidence that Gd accumulates in the brain and bones (Kanda *et al*., 2015; Lord *et al*., 2018), with unknown long-term effects. Finally, there is a growing concern for the associated mutagenic effects of Gd on coastal aquatic wildlife, particularly when exposed to ultraviolet radiation (Birka *et al*., 2016; Rogowska *et al*., 2018).

Gd is also limited in quantifying hemodynamic measures. The precise concentration time course of Gd in a compartment of blood following a bolus injection is difficult to determine using DSC MRI. This limits the ability to attain quantitative perfusion data, particularly when fitting a model to a signal time course (Willats and Calamante, 2012). In addition, Gd concentrations are confounded by recirculation and leakage from capillaries, which hinders bolus identification and perfusion quantification (Willats and Calamante, 2012).

It has long been known that deoxyhemoglobin (dOHb), the deoxygenated form of oxygen’s carrier protein in our blood, is paramagnetic (Pauling and Coryell, 1936). The dilution of dOHb through increases in blood flow results in an increase of MRI signal – this is the basis of functional MRI (fMRI), which localizes neuronal activity by measuring its effect on the blood oxygenation level-dependent (BOLD) effect (Kwong *et al*., 1992; Ogawa *et al*., 1990). The paramagnetic properties of dOHb were only recently exploited to generate a quantifiable vascular signal change in DSC MRI by transiently reducing the oxygen concentration of inhaled gas during a DSC scan and consequentially increasing the concentration of dOHb ([dOHb]) (Poublanc *et al*., 2021; Vu *et al*., 2021). Like a Gd bolus, a transient bolus of dOHb has been used to calculate *rCBV, rCBF*, and *MTT* (Poublanc *et al*., 2021; Vu *et al*., 2021). Perfusion measures from a dOHb bolus have also been correlated with those attained in arterial spin labeling for healthy and anemic subjects (Vu *et al*., 2021). Researchers have also quantified perfusion by inducing hyperoxic boluses, which result in a decrease to [dOHb] and increase of the subsequent MRI signal (MacDonald et al., 2018).

The optimal parameters for acquiring accurate and precise perfusion metrics using these contrast agents, including how calculated perfusion values compare when using Gd *vs* dOHb, are unknown. These parameters include the type of contrast agent, duration of the bolus, amount of contrast agent (in the case of dOHb, the hypoxic dosage (ΔS_a_O_2_)), baseline oxygen saturation (base-S_a_O_2_), and choice of reference voxel (i.e., artery or vein). To examine these potential dependencies, we applied a DSC MRI signal model that incorporates separate vessel compartments with both linear extravascular and quadratic intravascular relaxation rate contributions, as used in previous (f)MRI models (Uludag *et al*., 2009; Kjolby *et al*., 2006; Kjolby *et al*., 2009). This enables modeling of the tissue, arterial, and venous responses to generic contrast agents with arbitrary susceptibility values and bolus characteristics, such as duration and shape. We use this model to examine the effects of the aforementioned parameters on perfusion quantification. We validated the model by comparing the simulation predictions to the experimental results obtained from healthy subjects at 3T, utilizing dOHb boluses and a Gd injection in separate acquisitions.

## DSC MRI Signal Theory

### Section 1: Forward Model

The forward model is based on work by Uludag *et al*., 2009, originally developed to model gradient- (GRE) and spin-echo (SE) fMRI signals for magnetic field strengths between 1.5 and 16.4T (Duong *et al*., 2003; van Zijl *et al*., 1999). As the fMRI signal is caused by susceptibility changes induced by changes in dOHb, the model can also be applied to DSC MRI utilizing any contrast agent that induces magnetic susceptibility changes. Here, this framework is generalized and applied to model 3T GRE signal changes induced by boluses of both dOHb and Gd.

#### Signal Intensity

T_2_*-weighted MRI signal is composed of extravascular and intravascular signal contributions, weighted by their respective volumes (Uludag *et al*., 2009):

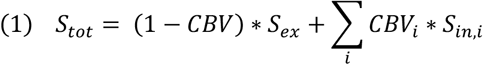

*S_ex_* and *S_in_* are the extravascular and intravascular signal contributions, respectively; both are weighted by the volume component where signal originates (*CBV*). The index *i* denotes the specific vascular compartment within the voxel being simulated (artery, vein, capillary, venule, or arteriole). Therefore, the signal is a summation of extravascular signal along with various intravascular signal contributions, each scaled by their volume proportion within the voxel. Of note, some DSC signal models from previous clinical research studies do not distinguish *S_in_* and *S_ex_* components (Calamante *et al*., 2000; Chappell *et al*., 2015; Patil *et al*., 2013; Patil and Johnson, 2011). The separate signal components can be written as:

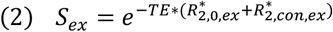

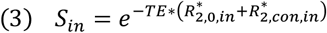

*S_ex_* and *S_in_* (Eqs. 2 and 3) are modeled as mono exponentials to the power of the negative echo time (TE is set to 30 ms) multiplied by the transverse relaxation rate 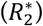, the inverse of the *T_2_*\* (Yablonskiy and Haacke, 1994). Here, 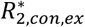 and 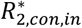 are the magnetic relaxation rates induced by applied contrast agent in the extravascular and intravascular space of a voxel, respectively. 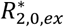 and 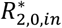 are the baseline magnetic relaxation rates for extravascular and intravascular space without any contrast agent present, respectively.

As shown in Eq. 1, separate signal exponentials are added together for each vascular compartment within a voxel for *S_in_*, however, relaxation rates are added from each vascular compartment within **a single** exponential for *S_ex_*, as will be described in Eq. 5.

#### Extravascular and Intravascular Relaxation Rates

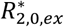 and 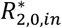 have been experimentally characterized for 3T at approximately 20.99 s^-1^ and 13.8 s^-1^, respectively for fully oxygenated blood (Zhao *et al*., 2007; Uludag *et al*., 2009). Polynomial equations that model 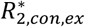 for various vessel radii have been determined using Monte-Carlo simulations – for a range of susceptibility values at 3T, the equations (expressed as per one percent of *CBV*) can be simplified by a linear relationship (see Table 2 in Uludag *et al*., 2009):

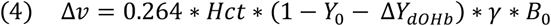

**Table 1.**
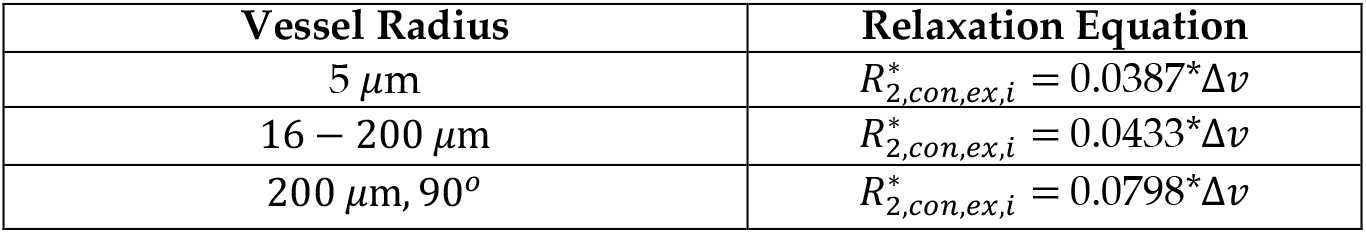
Coefficients for Extravascular Relaxation Rates in Various Vessel Radii. 5 *μm* (capillary), 16 – 200 *μm* (large artery and vein randomly oriented to the magnetic field, arteriole, and venule), and 200 *μm, 90°* (artery and vein perpendicular to the magnetic field) are the simulated vessel radii. Note that the large artery and vein can also be chosen to be randomly oriented, as is shown in Uludag *et al*., 2009, which affects the quantitative predictions but not the main findings (data not shown). *Δv* represents the frequency shift at the surface of the vessel in the presence of contrast agent, shown in Eq. 4.

**Table 2.** Summary of Simulated Voxel Properties for Arterial, Tissue, and Venous Voxels. The values in this table can be used for any arbitrary set of physiological assumptions. Vessel ratios and *ϒ_0_* for each vessel type were informed by previous work (Uludag *et al*, 2009; Vovenko, 1999). The arterial voxel is comprised of extravascular space and an artery that is oriented 90° to the main magnetic field, where the fraction of intravascular space is scalable from 0% to 100% (Eq. 1). Similarly, the venous voxel is comprised of extravascular space and a vein that is oriented 90° to the main magnetic field. These large vessels were simulated as perpendicular to the main magnetic field to mimic the stem of the middle cerebral artery (MCA) and dorsal portion of the superior sagittal sinus (SSS), which are both roughly perpendicular to the main magnetic field. *MTT* and delay, as shown in Table 2, were based on previous findings (Ibaraki *et al*., 2007).

|  | Arterial | Tissue | Venous |
| --- | --- | --- | --- |
| CBV (%) | 10-100 | 4 | 10-100 |
| MTT (s) | N.A. | 3 | 4 |
| Delay (s) | 0 | 1 | 2 |
| Vessel Composition | 90° Artery | Artery / Arteriole; Capillary; Vein / Venule (1:2:2) | 90° Vein |
| $Y_0$ (%) | 100 | Artery (100); Arteriole (95); Capillary (77.5); Vein/Venule (60) | 60 |

*Hct* represents the hematocrit fraction (simulated as 0.4), *Δϒ_dOHb_* represents the percentage change in the oxygenation as a decimal (which is caused by changes in dOHb), *γ* is the gyromagnetic ratio (2π * 42.6 MHz/T), and *B*_0_ is the magnetic field strength (3T). *ϒ_0_* represents the baseline oxygen saturation as a decimal, dependent on the vascular compartment (refer to Table 2). 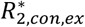 can then be found by summating 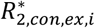 for each vessel compartment within the voxel:

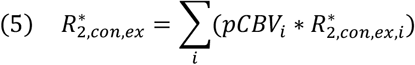

Here, *i* denotes an individual vessel compartment and *pCBV_i_* represents the percentage blood volume of the vessel contributing to the signal. Note that *pCBV_i_* must be a percentage and not a decimal (i.e., for a blood volume of 5%, a value of 5 is used as opposed to 0.05), as the equations for 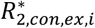 in Table 1 are modeled for one percent of *CBV*.

Finally, the 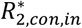 dependence on dOHb at 3T (*Hct* ≈ 0.4) was found to be quadratic at 3T from previous blood phantom work (Zhao *et al*., 2007):

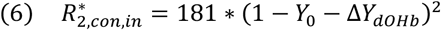

While quadratic modeling of the intravascular contrast relaxation rate is shown in numerous papers, some researchers have instead proposed a linear model to describe the intravascular relaxation rates (Lind *et al*., 2020; Wilson *et al*., 2017). The linear relationships were ruled out in our work (and later shown that the experimental results are not compatible with a linear relationship) – for Lind *et al*., this value was determined *in vivo* and likely suffered high partial volume errors; the value 89 mM^-1^s^-1^ is far more reflective of extravascular relaxivity (Lind *et al*., 2020; Kjolby *et al*., 2006). For Wilson *et al*., the concentration of contrast agent was studied at levels beyond those typically observed (Wilson *et al*., 2017). This will be elaborated on in the Discussion section.

#### Quantifying the Relationship Between dOHb and Gd

The above relationships allow for the quantification of the MRI signal in response to changes in dOHb, but to calculate the signal change induced by Gd, the effect of Gd on 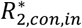 and 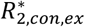 must be determined. One way to determine this is by finding a quantitative relationship between Gd and dOHb – given that dOHb and Gd are both paramagnetic contrast agents, a certain increase in [dOHb] will induce the same susceptibility effect as 1 mM of Gd.

The frequency shifts caused by changes in dOHb and Gd are both known. The frequency shift induced by dOHb is described in Eq. 4 while the frequency shift induced by Gd (when *Hct ≈* 0.4) is described by the following equation, derived from work in Kjolby *et al*., 2006:

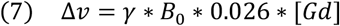

[Gd] represents the concentration of Gd (mM) in blood, and 0.026 ppm/mM represents the molar susceptibility of Gd, specifically Gd-DTPA (van Osch *et al*., 2003). Setting Eq. 7 equal to Eq. 4, assuming a *ϒ_0_* of 100%, allows for the interconversion of percentage dOHb saturation and Gd concentration (mM). This interconversion is a critical step in being able to simulate and compare dOHb and Gd-induced signal changes using the same DSC MRI framework:

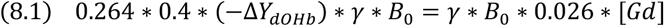

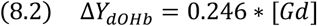

By setting *Δv* induced by dOHb equal to that induced by Gd, and setting the field strength to 3T, a certain concentration of Gd associated with magnetic susceptibility and 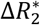 values can be converted to a value of blood oxygenation. For example, a 1 mM increase in Gd is equivalent in susceptibility to a 24.6% increase in dOHb at 3T, this is referred to as the pseudo-oxygenation (the oxygenation change that has the same effect as 1 mM of Gd on the magnetic field). Eqs. 1–6 were then used to simulate either a Gd- or dOHb-induced bolus by substituting *ΔY_d0Hb_* with 0.246 * [Gd], as described in Eq. 8.2. Thus, in our work, Gd and dOHb were modeled with the exact same signal framework, with differences only in the input bolus properties (susceptibility value, concentration, shape, and duration).

This can be further understood in Figure 1, which shows that while typical Gd and dOHb boluses occupy distinct peak susceptibility ranges, it is theoretically possible to increase [dOHb] or reduce [Gd] such that these contrasts agents would then have the same susceptibility range and therefore perfusion values quantified would be identical. This idea is further explored in the Discussion.

**Figure 1.**
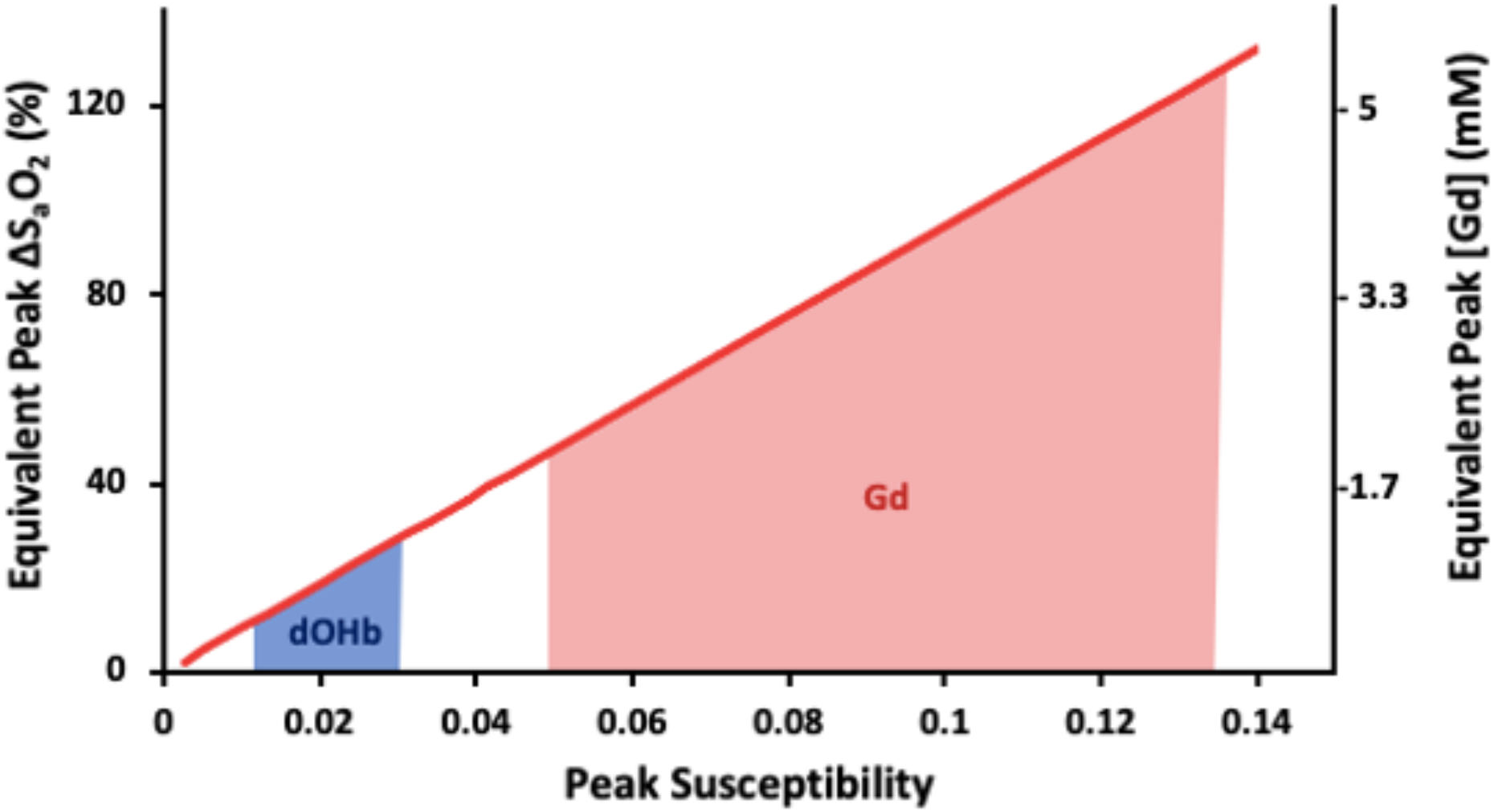
Susceptibility Relationship of Gd and dOHb in Simulations. Peak ΔS_a_O_2_ (%) and Gd (mM) as a function of peak susceptibility induced by contrast agent. The region shaded in blue is the typical peak susceptibility elicited in a simulated hypoxic bolus (Peak ΔS_a_O_2_ ≈ 10-25%). The region shaded in red is the typical peak susceptibility elicited in a simulated Gd bolus (Peak [Gd] ≈ 2-5 mM).

### Section 2: Model Inversion

#### Relaxation Curves

The first step in standard DSC analysis, following preprocessing, is conversion of the signal time course (*S*(*t*)) into the relaxation rate time course 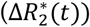:

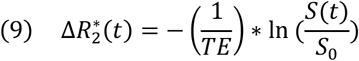

This step requires the specification of a baseline signal (S_0_), prior to the administration of contrast agent, along with the *TE*.

#### Cerebral Blood Volume

From this point forward, the term ‘relative’ will be used to describe tissue signal normalized by signal from a reference voxel (arterial or venous). Once tissue, arterial, and venous signals are converted into their respective 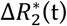 curves, the relative *CBV* (*rCBV*) can be calculated:

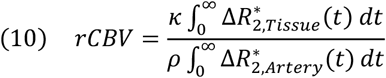

Here, by taking a ratio of the tissue integral to the arterial integral, tissue *rCBV* can be estimated. We also examine the effect of replacing 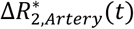 with 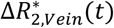. Normalizing to venous as opposed to arterial signal is of clinical interest, given that venous voxels tend to have reduced partial volume effects (Calamante *et al*., 2013).

*ρ* represents the brain density (1.05 g/mL) and κ represents the hematocrit correction for micro- and macrovasculature ((1 – *Hct*)/(1 – 0.69*Hct*)) in Gd contrast (Tudorica *et al*., 2002). In dOHb contrast, we concluded that the κ value should theoretically be set to *Hct*/0.69*Hct*, as contrast is limited to the red blood cell fraction and not the plasma. These scaling factors are implemented in the analysis of the experimental data but not included in the numerical simulations (neither in the forward model nor model inversion).

#### Cerebral Blood Flow and Mean Transit Time

In DSC MRI analysis, 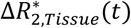 is assumed to be equal to the convolution of 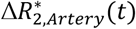 and the tissue residue function (*R*(*t*)), further scaled by the relative *CBF* (*rCBF*). *R(t*) represents the dispersion of contrast agent as it passes through the vasculature of a tissue voxel. This equivalency is based on the indicator dilution theory and is described in Eq. 11 (Meier and Zierler, 1954).

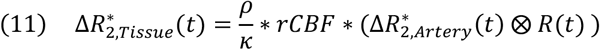

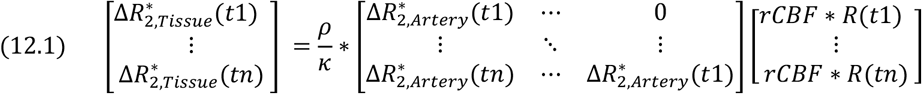

For our simulations, the convolution is discretized as illustrated in Eq. 12.1 (Ostergaard *et al*., 1996; Wirestam *et al*., 2000). This allows for the deconvolution of 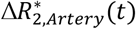 from *R*(*t*) using singular value decomposition (SVD), a popular technique in DSC MRI analysis (see Willats and Calamante, 2012).

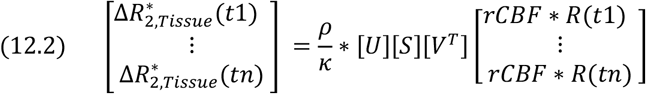

In this method, the arterial matrix is broken down into the product of *U*, *S* (singular values), and *V^T^* (Eq. 12.2) – these resulting matrices are rearranged to solve for *R*(*t*), which is scaled by *rCBF*, taken as the maximum height of *R*(*t*) (Ostergaard *et al*., 1996; Wirestam *et al*., 2000). Although no noise is added in the simulations, it is present in the experimental data. Therefore, we also explored using a threshold in both the simulations and experimental validations to remove singular value components less than 20% of the maximum singular value in *S*. A threshold of 20% was previously determined to provide the most accurate estimations of *rCBF*, given typical noise levels of experimental data (Bjornerud and Emblem, 2010).

Finally, *MTT* (in seconds) is calculated as a ratio of *rCBV* to *rCBF*, which follows from the central volume theorem (Meier and Zierler, 1954):

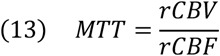

## Methods

### Part 1: Simulation Methods

#### Arterial, Tissue, and Venous Concentration Input Functions

The Gd input time course (Gd(t)) in artery is well-modeled as a gamma variate function (Davenport, 1983; Thompson *et al*., 1964).

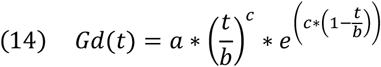

While it is difficult to attain the exact concentration profile of Gd in an artery (Kellner *et al*, 2018; Lind *et al*, 2020; Patil *et al*., 2013), the shape parameters, *b* (time to peak) and *c*, can be set to 2.5 s and 3 (unitless) respectively, to roughly model the shape observed experimentally. *a* (the peak concentration) can be set to approximately 2 mM – see Discussion. The dOHb input, as dictated by our experimental validations, and previous work (Poublanc *et al*., 2021), was modeled as a rectangular hypoxic bolus of roughly 30 s duration, convolved with a decaying mono-exponential as the bolus takes some time to reach the target oxygenation and enter the brain. Note that the forward model, as described above, can be used for any shape, duration, and concentration for both Gd and dOHb inputs.

To attain the tissue concentration profile, the arterial input was convolved with a well-defined biexponential residue function (*R*(*t*)) which designates fast and slow flowing vascular compartments (Mehndiratta *et al*., 2014):

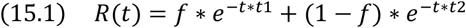

Here *t1* (fast-flowing compartment transit time), *t2* (slow-flowing compartment transit time), and *f* (fraction of flow in the fast-flowing compartment), are vascular flow parameters, which yield the *MTT* in the following equation (Mehndiratta *et al*., 2014):

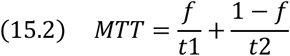

As we expect the venous voxel to be largely composed of a single vascular compartment, we convolve the arterial input with a mono-exponential residue function to obtain the venous concentration profile (Mehndiratta *et al*., 2014):

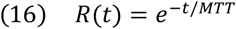

The flow parameters, informed by clinical data and literature yield a tissue voxel *MTT* of roughly 3 s (*f* = 0.92, *t_1_* = 0.68, *t_2_* = 0.05) and a venous voxel *MTT* of 4 s (*MTT* = 4) (Ibaraki *et al*., 2007). As well, a delay of 1 s and 2 s were applied to the tissue and venous voxels, respectively (Ibaraki *et al*., 2007). These parameters are easily adjustable to model other values.

#### Simulated Voxel Parameters

### Part 2: Experimental Methods

#### Subjects

This study was approved by the Research Ethics Board of the University Health Network, Toronto, Canada, and conforms to the Declaration of Helsinki. Written informed consent was obtained in all 6 healthy volunteers (age range 22-60, 1 woman). All subjects were non-smokers and not taking regular medication.

#### Scan Protocols and MRI Sequences

The RespirAct™ RA-MR (Thornhill Medical, Toronto, Canada) was used to target lung S_a_O_2_ while maintaining isocapnia (Sobczyk *et al*., 2016). All gases and sensors were calibrated prior to use. GRE BOLD and anatomical T1 scans were acquired on a 3T MRI system (Signa HDx - GE Healthcare, Milwaukee) with echo-planar (EPI) acquisition.

MR parameters for dOHb and Gd GRE BOLD scans were TR/TE = 1500/30 ms, Flip angle = 73°, 3 mm isotropic voxels, 29 contiguous slices – no interslice gap. dOHb scan duration was approximately 210 s per paradigm; Gd scan duration was approximately 20 s. A high-resolution T1-weighted 3D spoiled GRE sequence was acquired for each subject as well (TI = 450 ms, TR = 7.88 ms, TE = 3 ms, flip angle = 12°, voxel size = 0.859 x 0.859 x 1 mm, matrix size = 256 x 256, 146 slices, field of view = 24 x 24 cm, no interslice gap).

For the Gd paradigm, 5 mL of gadobutrol (Gadovist®) was injected at 5 mL/sec followed by 30 mL of saline at 5 mL/sec.

In the dOHb experiments, hypoxic dosage (ΔS_a_O_2_) and/or baseline oxygen saturation (base-S_a_O_2_) were varied (Figure 3). To determine the effect of ΔS_a_O_2_ and base-S_a_O_2_ for perfusion quantification, S_a_O_2_ values were modified to generate four hypoxic paradigms with three ~30 s hypoxic boluses as follows: 98% to 90%, 98% to 84%, 98% to 75%, and 88% to 80%. Each hypoxic drop was acquired over multiple breaths, depending on rate and volume of ventilation, as attaining the hypoxic state required wash-out of oxygen in the functional residual capacity of the lungs (air remaining after exhalation). Thus, the duration of the dOHb bolus was longer than for Gd, which in contrast depends on the speed of injection and dispersion of the bolus in its trajectory from peripheral vein to the brain. The hypoxic target was maintained for approximately 30 s before returning to baseline in a single breath. The rapidity of the return to baseline was due to the higher available oxygen partial pressure target (713 mmHg) relative to the hypoxic target (<100 mmHg). The baseline before and after each bolus was 30 s in duration.

**Figure 2.**
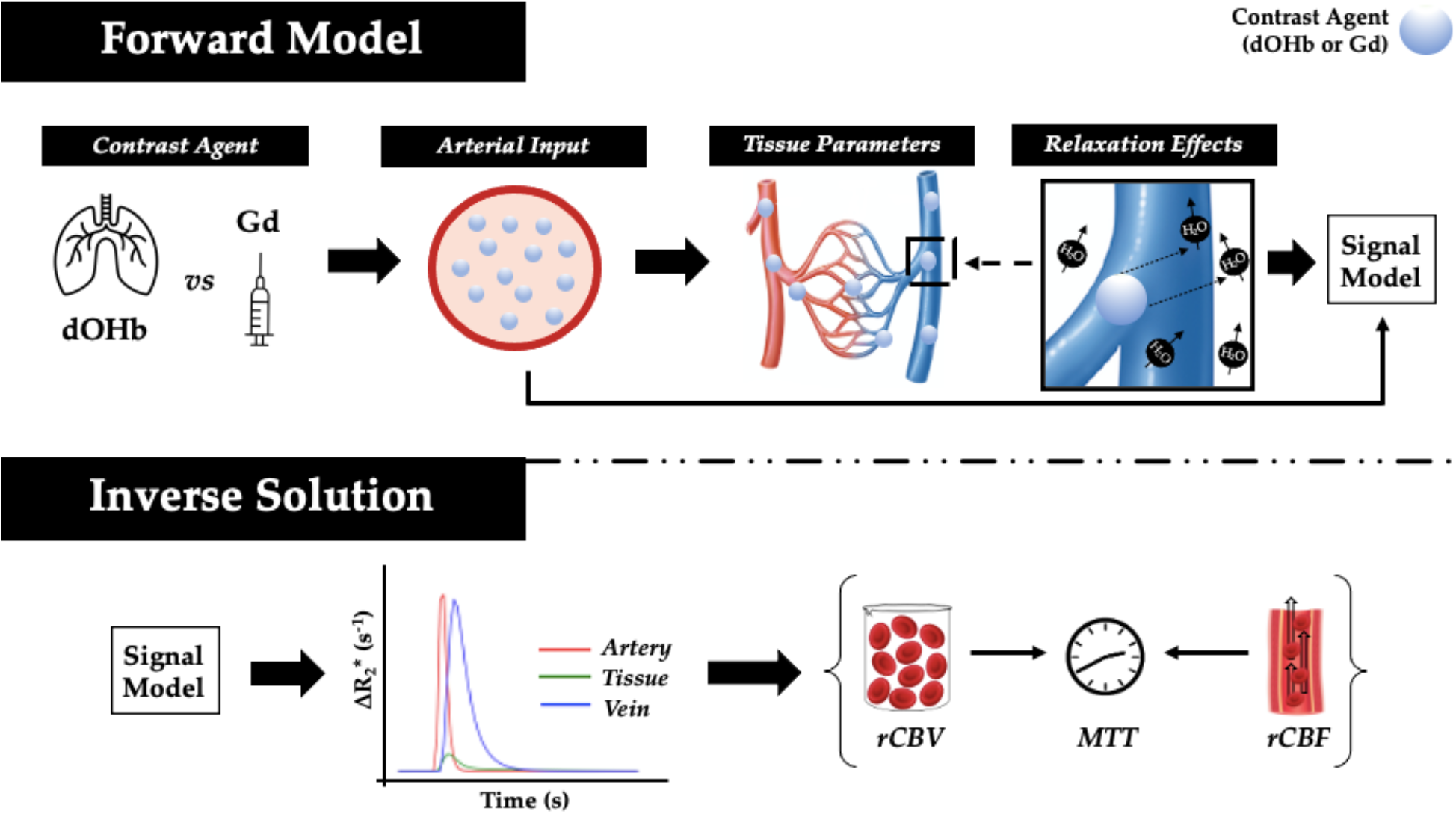
Summary of Simulation Method. **A. Forward Model (top).** Contrast profiles for Gd [mM] and dOHb (S_a_O_2_ (%)) are simulated; tissue, arterial, and venous properties (*CBV*, *MTT*, *CBF*) are then applied; the *ΔR_2_*_ex_* and *ΔR_2_*_in_* response for water protons are modeled; finally, the resultant 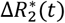 time courses are fed into a signal model. **B. Inverse Solution (bottom)** Tissue, arterial, and venous *S(t*) curves are generated from Eq. 1. These curves are converted to 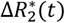 (Eqs. 9). The arterial input and tissue response curves are then used to calculate *rCBV* (Eq. 10), *rCBF* (Eqs. 11–12), and the *MTT* (Eq. 13).

**Figure 3.**
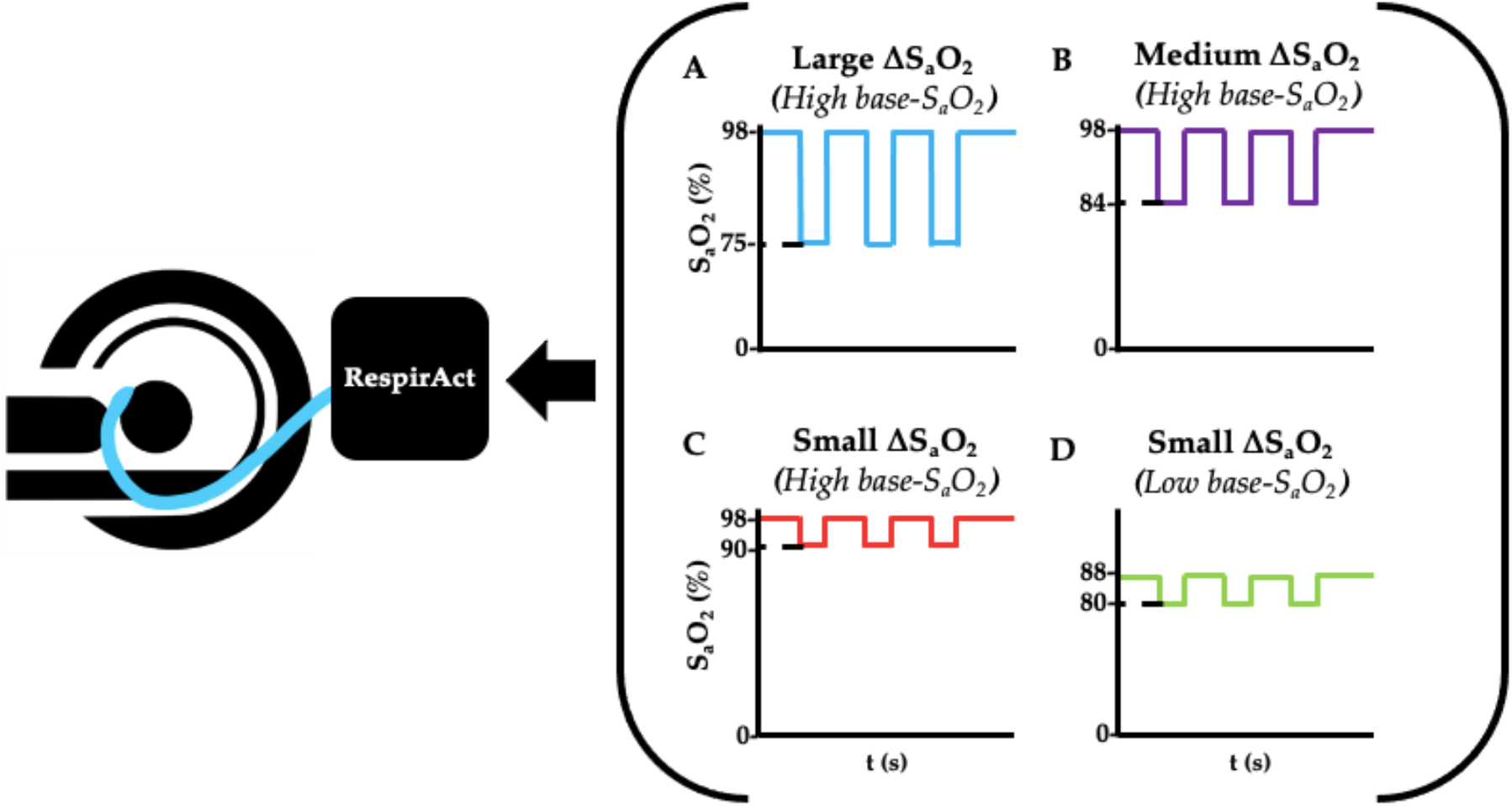
Summary of the dOHb Experimental Method. Subjects (n=6) were imaged in a 3T MR scanner with a mask attached to the RespirAct®, where S_a_O_2_ was altered to attain desired levels of base-S_a_O_2_ and ΔS_a_O_2_. In sum, the following paradigms were generated: small ΔS_a_O_2_ with high base-S_a_O_2_ (red: 98% to 90% S_a_O_2_), medium ΔS_a_O_2_ with high base-S_a_O_2_ (purple: 98% to 84% S_a_O_2_), large ΔS_a_O_2_ with high base-S_a_O_2_ (blue: 98% to 75% S_a_O_2_), and small ΔS_a_O_2_ with low base-S_a_O_2_ (green: 88% to 80% S_a_O_2_).

One subject reported a mild change in respiratory sensation during the dOHb protocol. Two subjects reported discomfort and nausea during Gd acquisitions. Only data from one paradigm of one subject was removed due to excessive motion during the scan.

#### Preprocessing

FSL (version 6.0.4) was used for image preprocessing (Woolrich *et al*., 2009): brain extraction was performed on T_1_ anatomical images; brain extraction, motion correction, interleaved slice time correction, and Gaussian temporal filtering were performed on BOLD scans. Gaussian temporal filtering was only performed on the dOHb BOLD scans by averaging each signal time point by a 1×5 Gaussian kernel. For all BOLD scans, the first 2-3 timepoints were removed to allow for equilibration. BOLD signal and SNR were calculated according to:

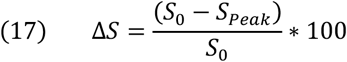

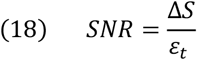

In Eq. 17, *ΔS* is the percentage maximal signal change, S_0_ is the average signal of ten temporal volumes before and after the selected bolus, and *S_Peak_* is the maximum signal drop. The signal to noise ratio (*SNR*) is computed as shown in Eq. 18, where *ε_t_* is the standard deviation of signal for ten temporal volumes before and after the selected bolus divided by *S*_0_. *S(t*) was then converted to 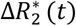 using Eq. 9. The above equations were applied to dOHb and Gd signal data from all subjects using Python 3.7.6.

#### Selection of Arterial and Venous Voxels

To reduce registration steps and, hence, misregistration errors, we sought to identify arterial and venous voxels in the functional rather than the anatomical space. As arterial voxels are difficult to clearly identify on functional images, we used the following procedure for selection of the voxels: Arterial voxels were defined based on their proximity to the M1 segment of the MCA using registered T_1_ anatomical data, *ΔS* within the top 10% of the *ΔS* values, and short delay (~ 0-1 s). Venous voxels were defined based on their proximity to the SSS using registered T_1_ anatomical data, *ΔS* within the top 10% of the *ΔS* values, long delay on the Gd and dOHb maps (~ 2 s), and low *S_0_* (the reason for a low *S_0_* is that vein consists of a higher dOHb saturation at rest, which reduces the overall signal). Once the arterial or venous voxel of interest was selected on the Gd map, the selected voxel was linearly registered to each dOHb paradigm to avoid displacement of the voxel between the different paradigms.

#### Perfusion Quantification

After defining an arterial input function (AIF) from the M1 segment of the MCA or venous output function (VOF) from the dorsal section of the SSS, each time series was truncated to begin at the start of the AIF bolus and finish at the end of the VOF bolus. Then, *rCBV*, *rCBF*, and *MTT* for each voxel were calculated using 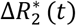 and Eqs. 10–13. Any voxel yielding a negative value for *rCBV, rCBF, or MTT*, was excluded from further analysis – this was observed in voxels with high noise and low blood volume (more often for dOHb than Gd).

#### Grey and White Matter Segmentation

The *ΔS* functional maps for Gd were segmented into grey matter (GM) and white matter (WM) using a threshold method. WM voxels were defined as those containing *ΔS* values within the lowest 15-20% of the *ΔS_G_d* data range, while GM was defined as voxels containing the highest 75% *ΔS* values. The maps were then binarized and linearly registered to the functional space of each dOHb paradigm. The GM and WM regions were further masked with a thresholded *ε_t_* map (voxels with values in the lowest 10% of the *ε_t_* range were kept in mask) to remove noisy voxels in peripheral brain tissue and skull base. The exclusion of voxels based on *ε_t_* also removes large arterial and venous voxels from the GM masks due to high levels of physiological noise in these regions. The GM and WM were finally masked with a thresholded Gd *MTT* map (voxels within the lowest 70% of robust range of *MTT* were kept in the mask) to remove voxels in the CSF (Figure S2).

#### Data and Code Availability

Anonymized data will be shared by request from any qualified investigator for purposes such as replicating procedures and results presented in the article, so long as the data transfer is in agreement with the University Health Network and Health Canada legislation on the general data protection regulation.

Code will be made available via GitHub upon acceptance of paper.

## Results

Figure 4 displays *S(t*) and 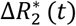 for simulated tissue, arterial, and venous voxels. The baseline signal is highest for artery and lowest for vein due to differences in baseline oxygen saturation. In addition, the dOHb boluses are more sluggish as compared to Gd boluses, mimicking experimentally observed bolus durations (roughly 30-40 s for hypoxic boluses). According to this simulation, 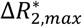, the peak of 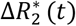, attained with dOHb is roughly 4 times larger in vein relative to artery even though the same ΔS_a_O_2_ is simulated for artery and vein, indicating a substantial effect of the baseline oxygen saturation on 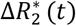. Remarkably, for Gd, 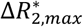 is only roughly 1.1 times larger in vein relative to artery, indicating a reduced effect of the baseline oxygen saturation on 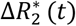 for contrast agents eliciting high susceptibility changes. As well, the ratio of arterial to tissue 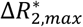 is much higher when using Gd in comparison with dOHb. Conversely, the ratio of venous to tissue 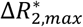 is much more consistent when using Gd versus dOHb.

**Figure 4.**
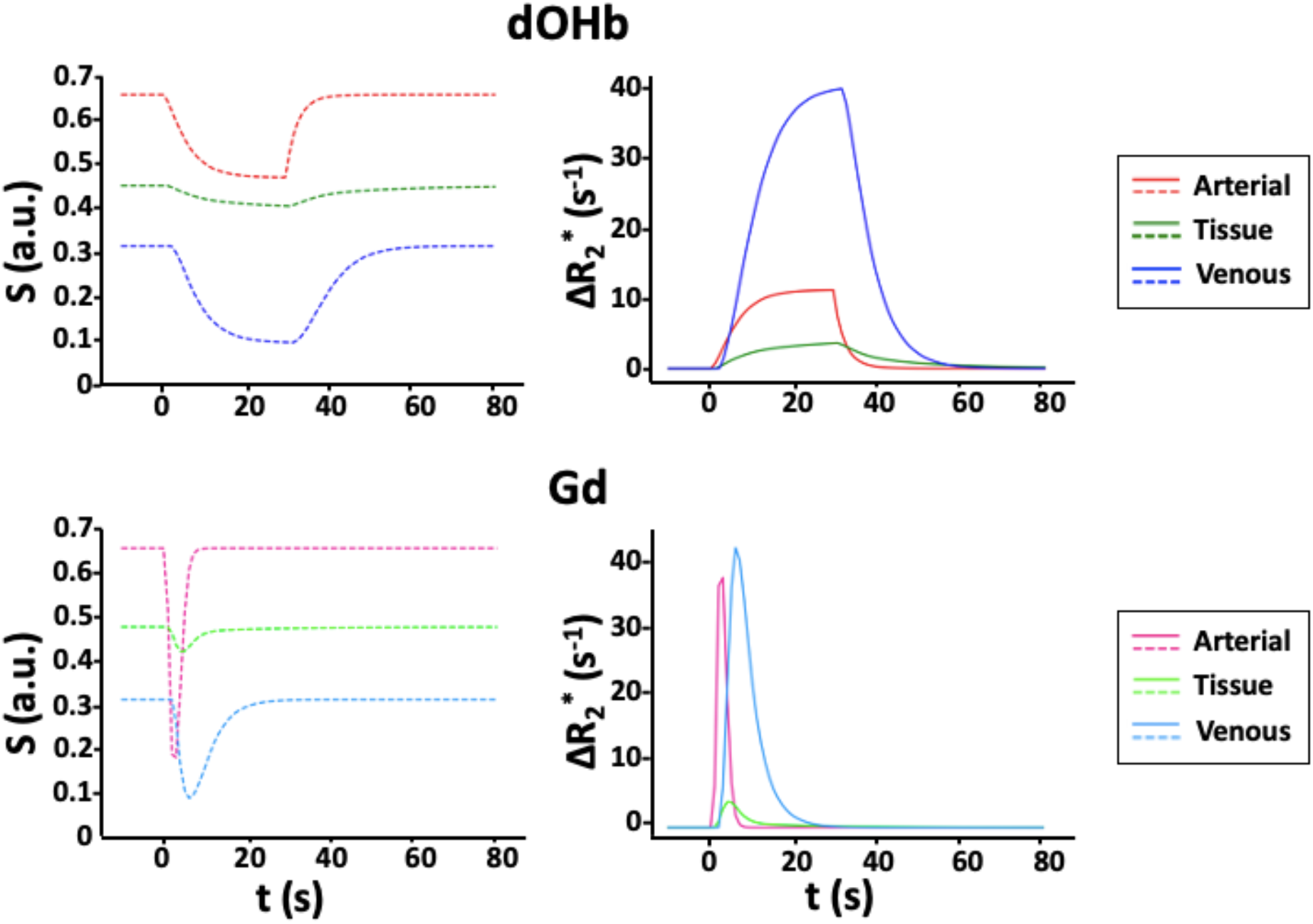
Simulated *S(t*) and 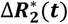 for Gd and dOHb Contrast. **Left.** *S(t*) curves in artery (*CBV* = 100%), tissue (*CBV* = 4%, *MTT* = 3 s), and vein (*CBV* = 100%, *MTT* = 4 s) following contrast administration of dOHb (top) and Gd (bottom). **Right.** Corresponding 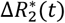 curves. Note that the simulated dOHb profile follows a 98% to 75% S_a_O_2_ and Gd profile has a peak input concentration of 2 mM.

Figure 5 provides a more detailed analysis of the 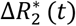 curves in artery, tissue, and vein at multiple simulated *CBV*s. For tissue, linear changes in the simulated *CBV* result in increased changes in the 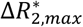. This is not the case for arterial and venous responses. For the arterial response to dOHb, increasing *CBV* increases the 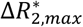 up to a certain CBV values, such that any further increase in *CBV* leads to decreases in 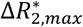. The simulated arterial *CBV* to 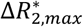 relationship in response to Gd has also a maximum for an intermediate CBV value. For venous voxels, 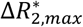 saturates for high CBV values for both Gd and dOHb. These results point to a complex relationship between relaxation rate changes and tissue composition.

**Figure 5.**
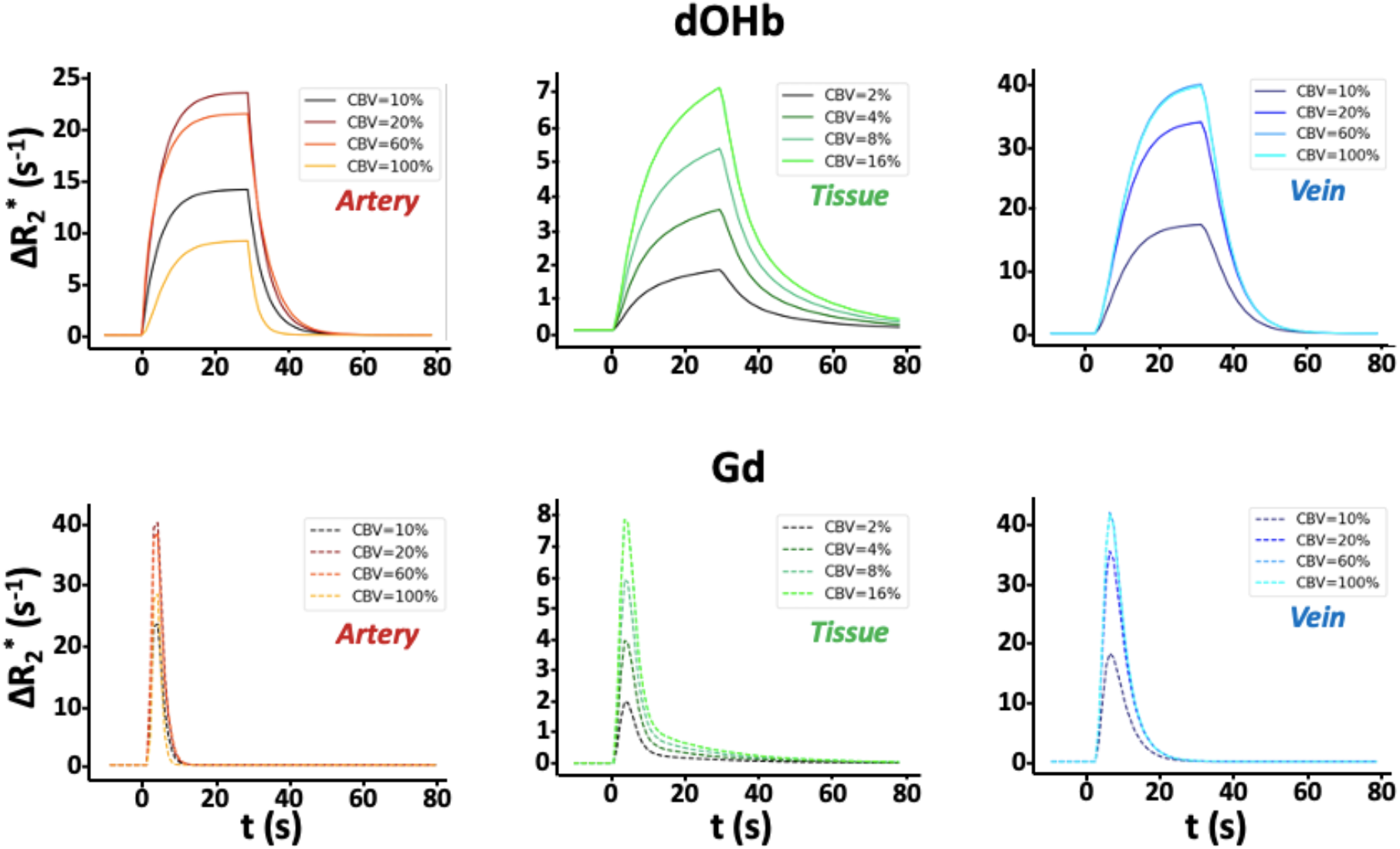
Simulated 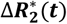 for dOHb and Gd Contrast. **Left.** 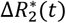 for arterial voxel with varying intravascular space in response to dOHb (top) and Gd (bottom). **Middle.** 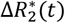 for tissue voxel with varying intravascular space in response to dOHb (top) and Gd (bottom). **Right.** 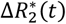 for venous voxel with varying intravascular space in response to dOHb (top) and Gd (bottom). Note that tissue curves (*MTT* = 3 s) are convolved with a more quickly decaying exponential than venous curves (*MTT* = 4 s).

Figure 6 shows the comparison of *ΔS* acquired with dOHb and Gd boluses. For simplicity here and elsewhere, *ΔS* is displayed as an absolute signal change. *ΔS_Gd_* is ~10x higher than *ΔSdOHb*. When the *ΔSdOHb* values are divided by the *ΔS_G_d* values in corresponding voxels, hereafter termed the *ΔS* ratio, differences between large vessels, CSF, GM, and WM are visually highlighted (see Figure S2). The ratio of GM to WM with Gd is slightly larger than that of dOHb (roughly 2.6 for Gd and 1.9 for dOHb). The *ΔS* ratio in large vessels (artery and vein) is much higher than in GM or WM, as evident from inspection of the figures and in the box and whisker plot of Figure 6. The experimental finding of the *ΔS* ratio being higher in large vessel voxels in comparison to tissue voxels is corroborated by simulation findings (Figure S4) – here, the average of the arterial and venous *ΔS* ratios is larger than either GM or WM *ΔS* ratio.

**Figure 6.**
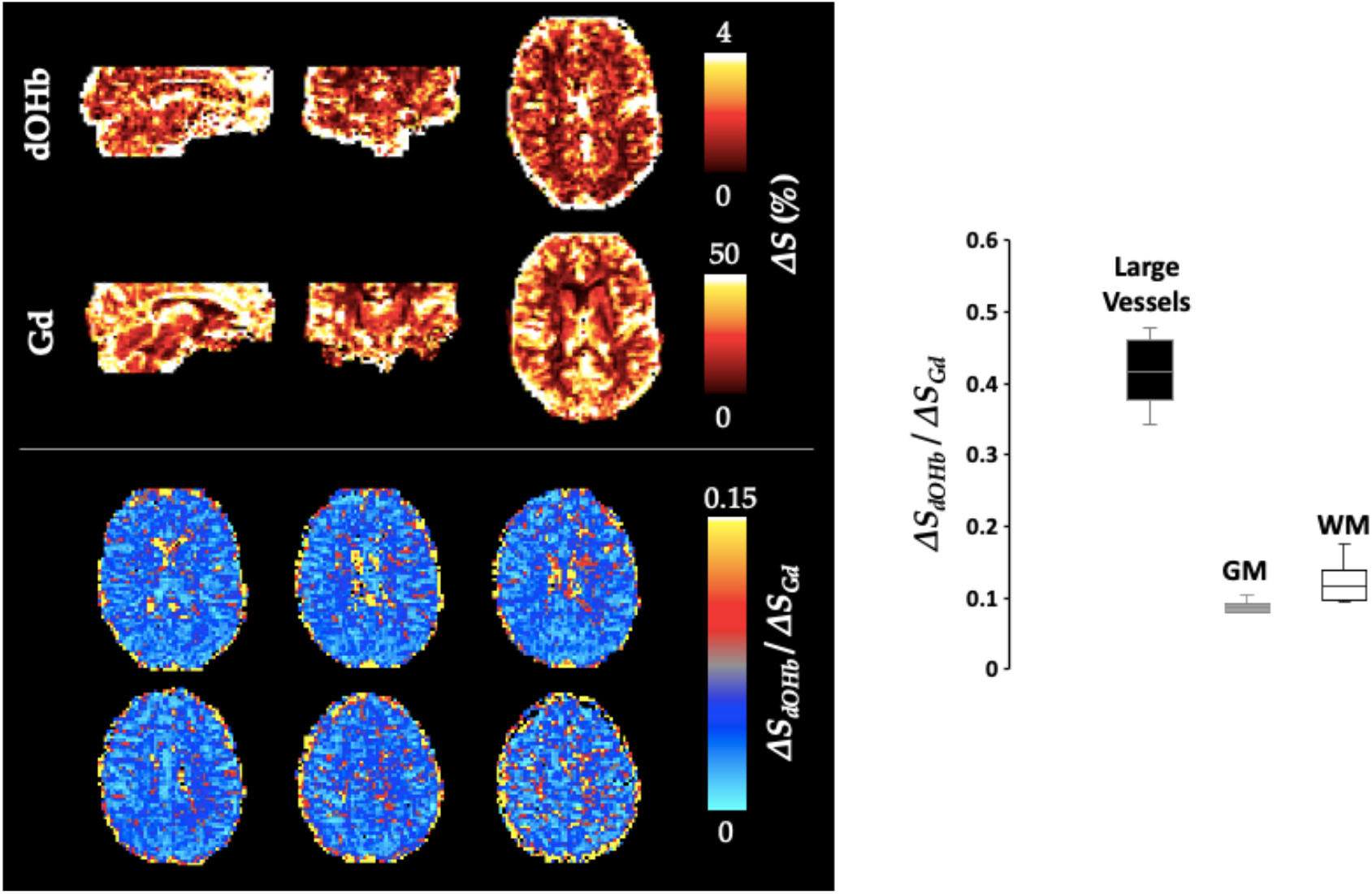
*ΔS* for Gd and dOHb in Vessels, GM, and WM. Left (Individual). *ΔS* (%) maps for Gd and dOHb in a representative subject (top). Note that dOHb map is an average of the following paradigms: small ΔS_a_O_2_ with high base-S_a_O_2_, medium ΔS_a_O_2_ with high base-S_a_O_2_, large ΔS_a_O_2_ with high base-S_a_O_2_. Six representative axial slices of Δ*S_dOHb_* map divided by *ΔS_Gd_* map (bottom). **Right (Summary).** Box and whisker plot for *ΔS_dOHb_/ΔS_Gd_* for various tissue compartments, including large vessels, GM, and WM. GM and WM masks described in methods; large vessel mask generated by taking highest 50% of *ΔS_Gd_* or *ΔS_dOHb_* robust range (n=6).

Figure 7 shows calculated *rCBV* maps from Gd and all dOHb paradigms in a representative subject when normalizing the tissue relaxation rate time course map to that of a selected arterial (top) or venous voxel (bottom) – refer to Eq. 10. We found that *rCBV* values depend on the extent of susceptibility change (contrast type (i.e., Gd or dOHb) and dOHb paradigm) when normalizing tissue to artery and vein, with a larger observed dependency on these contrast agent parameters when normalizing to artery. It is evident from the experimental validations that the order of *rCBV* estimation, from lowest to highest, is as follows: small ΔS_a_O_2_ with high base-S_a_O_2_, medium ΔS_a_O_2_ with high base-S_a_O_2_, large ΔS_a_O_2_ with high base-S_a_O_2_, and small ΔS_a_O_2_ with low base-S_a_O_2_. Thus, either increasing the ΔS_a_O_2_ or decreasing the base-S_a_O_2_ reduces the calculated *rCBV* values and can yield values more like those obtained from Gd data. The pattern of *rCBV* estimation for different paradigms when normalizing tissue to artery was consistent in all subjects, and the averages for each paradigm are shown in Table S1. When normalizing to vein, the pattern was consistent in 5 out of 6 subjects. Of note, the ratio of *rCBV_GM_* to *rCBV_WM_* is higher when using Gd in comparison to dOHb – this follows from the *ΔS* trend described in Figure 6.

**Figure 7.**
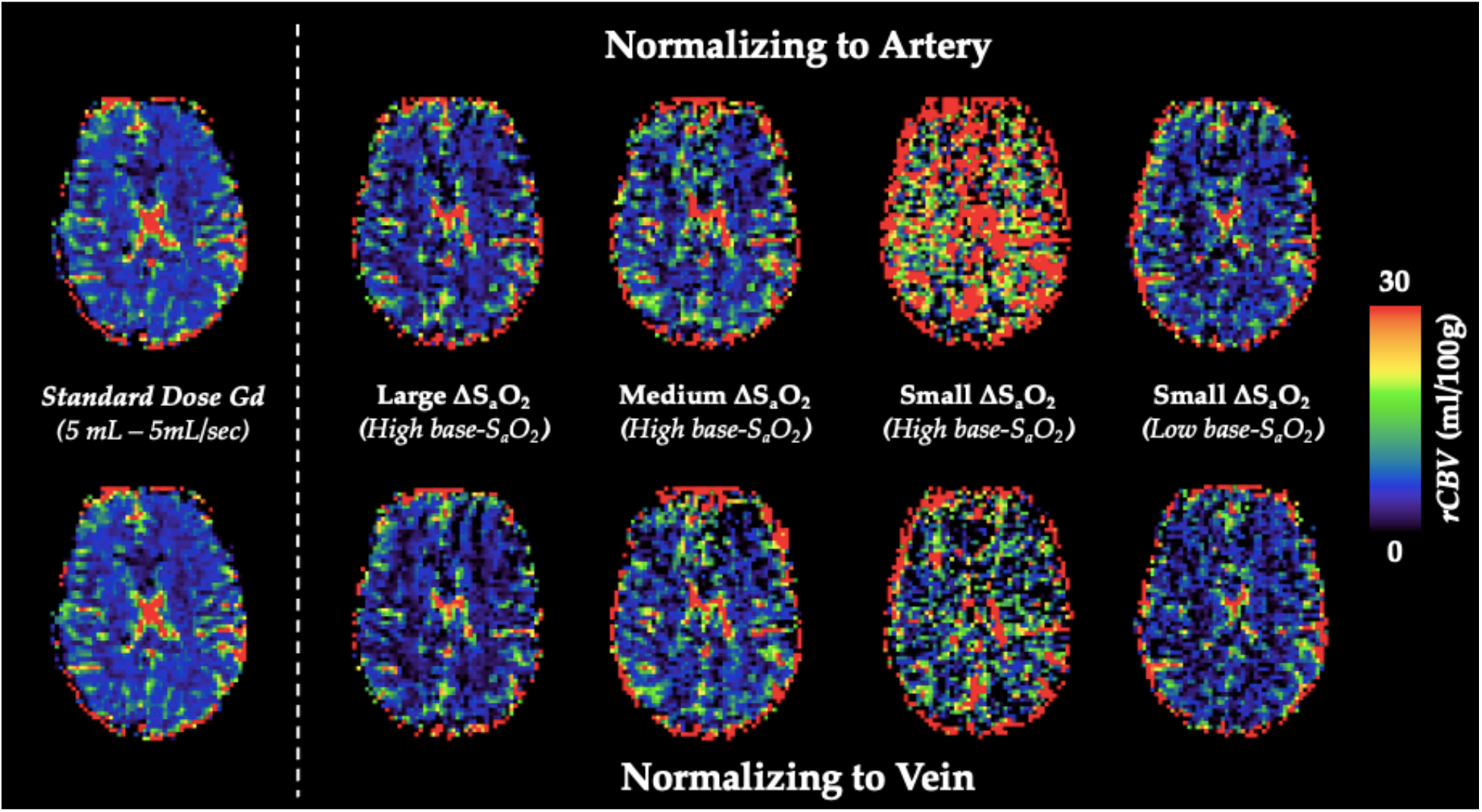
*rCBV* Dependency on ΔS_a_O_2_ and Base-S_a_O_2_. **Top (Arterial Normalization).** *rCBV* (mL/100g) maps from representative subject axial slice where tissue is normalized to an arterial voxel in the MCA for five subsequent paradigms (left to right): standard Gd, large ΔS_a_O_2_ with high base-S_a_O_2_, medium ΔS_a_O_2_ with high base-S_a_O_2_, small ΔS_a_O_2_ with high base-S_a_O_2_, small ΔS_a_O_2_ with low base-S_a_O_2_. **Bottom (Venous Normalization).** *rCBV* (mL/100g) maps from representative subject axial slice where tissue is normalized to a venous voxel in the SSS for the same paradigms.

Figure 8 displays *rCBV* results from both simulation and experimental data for Gd and all dOHb paradigms when normalizing tissue to artery versus vein. Here, the box and whisker plots show the calculated *rCBV* as whole brain (GM+WM) averages. When normalizing to artery, the trend is such that increasing the ΔS_a_O_2_ or decreasing the base-S_a_O_2_ reduces the magnitude of *rCBV*, making it more similar to that obtained when using Gd. The trend of *rCBV* estimations when normalizing tissue to vein is the same, but the effects of ΔS_a_O_2_, base-S_a_O_2_, and choice of contrast agent are lower in magnitude for both the simulations and experimental data. As well, using the vein to normalize tissue data for dOHb results in a much more comparable estimation of *rCBV* to both Gd estimation and the ground truth (Figure S1). Furthermore, as shown in the experimental validations, inter-subject variability (defined as the interquartile range in the box and whisker plot) is greatly reduced as the ΔS_a_O_2_ increases and base-S_a_O_2_ decreases (Figure 8). Ultimately, the experimental and simulation data show that normalizing tissue to vein as opposed to artery will result in more accuracy and consistency in the quantification of *rCBV* when using different dOHb paradigms.

**Figure 8.**
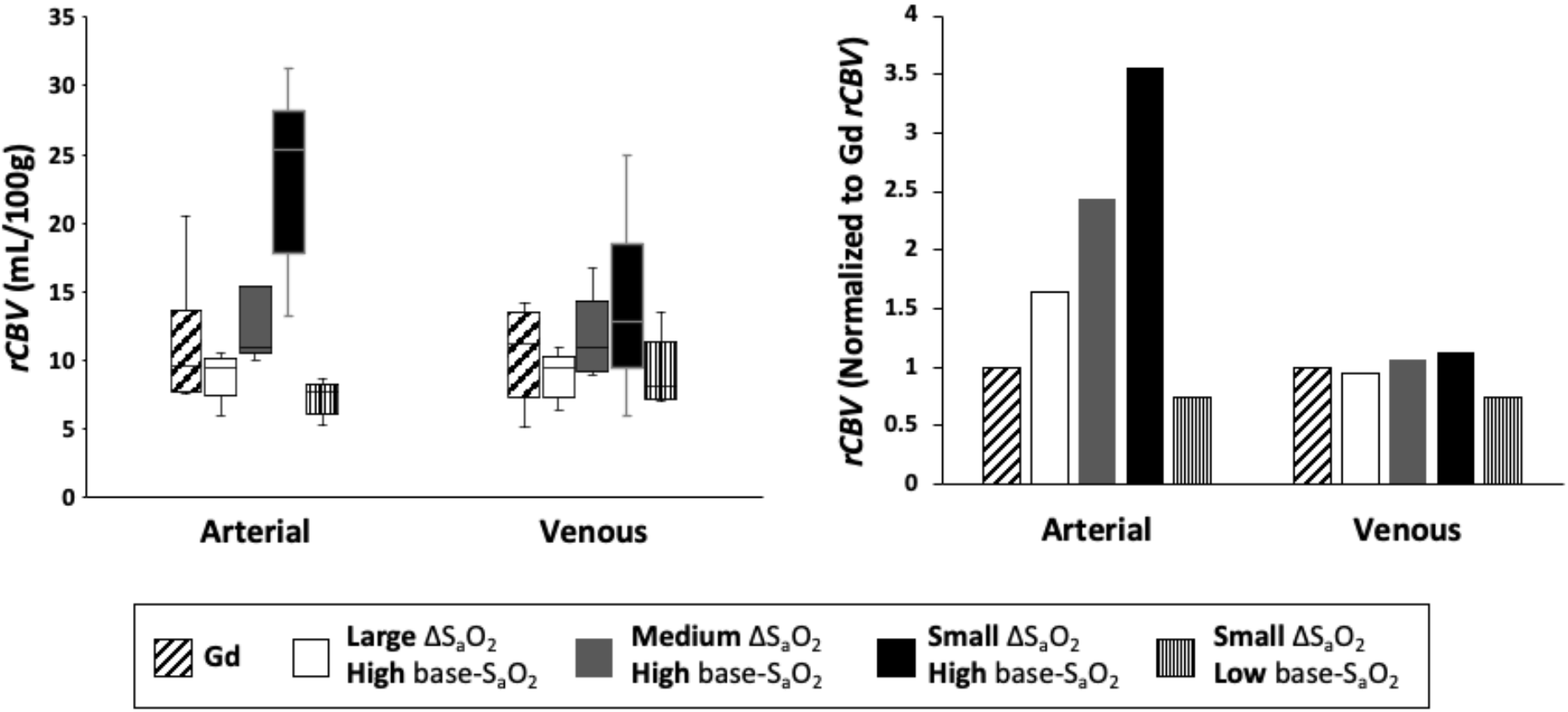
*rCBV* Dependency on Arterial *vs* Venous Normalization. **Left (Validations).** Box and whisker plot of calculated *rCBV* when normalizing tissue to arterial voxel in the MCA *vs* venous voxel in the SSS (n = 6) for Gd and dOHb at varying ΔS_a_O_2_ and Base-S_a_O_2_. **Right (Simulations).** Calculated *rCBV* when normalizing tissue to simulated artery *vs* vein for Gd and dOHb at varying ΔS_a_O_2_ and Base-S_a_O_2_. *rCBV* for all paradigms is further normalized to the *rCBV* attained with Gd (values above one are an overestimation relative to Gd). Simulated tissue *CBV* = 4%; Simulated arterial *CBV* = 100%; Simulated venous *CBV* = 100%.

Figure 9 displays simulation and experimental results for *MTT* attained with dOHb and Gd. For the experimental data, the estimated *MTT* values were greater when using dOHb than when using Gd contrast. This was observed in all dOHb paradigms for all subjects, as displayed in Table S1. While absolute values of calculated *MTT* differed in GM and WM between Gd and dOHb, the GM/WM ratio was generally consistent. The simulations show the same pattern as the experimental data, with dOHb contrast resulting in higher calculated *MTT* than Gd. We noted that this is especially the case when the singular values are thresholded in deconvolution, a necessary step in removing noise prior to *rCBF* and *MTT* estimation as described in the Methods. When thresholded, a dOHb bolus of longer duration results in a higher *MTT* estimation than a dOHb or Gd bolus of shorter duration. In sum, thresholding the singular values for deconvolution, a necessary step in the experimental data analysis, and using a bolus of longer duration both result in a greater overestimation of the *MTT*.

**Figure 9.**
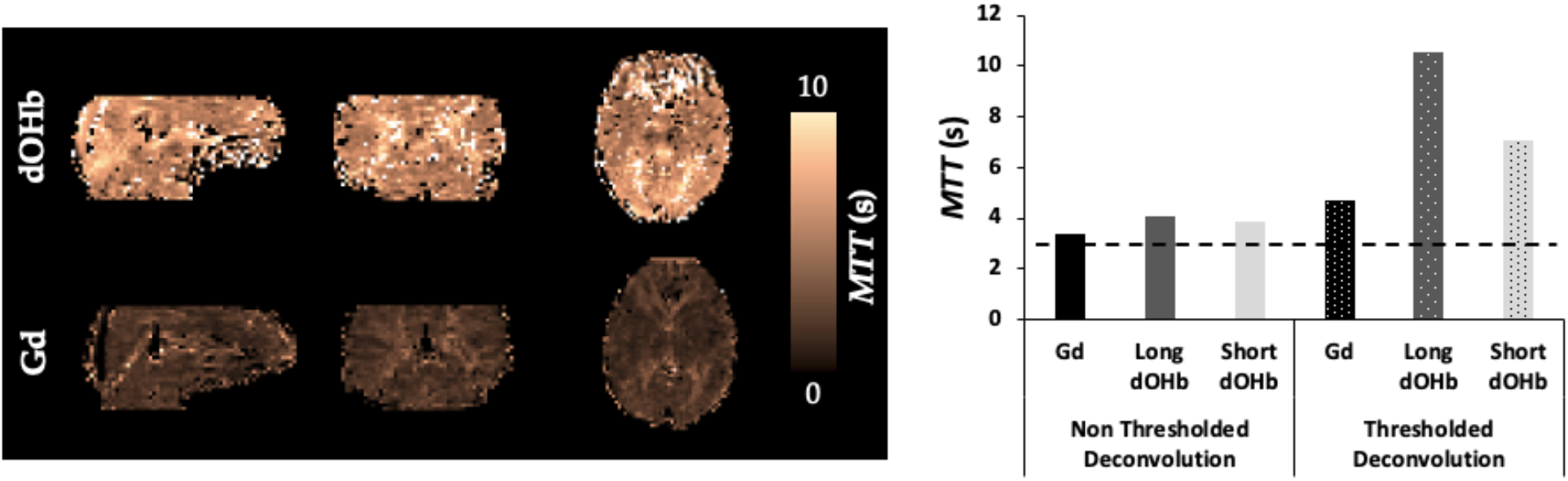
*MTT* Dependency on Bolus Duration. Left (Validations). *MTT* maps from a representative subject when exposed to a hypoxic bolus (dOHb) of roughly 30 second duration versus a Gd bolus of roughly 10 second duration. **Right (Simulations).** Calculated *MTT* with either no thresholding (solid) or standard deconvolution thresholding (dotted) for dOHb and Gd contrast (no noise). Simulated dOHb bolus is either longer (dark gray) or shorter (light gray) to match the duration of a standard Gd bolus (black). The ground truth *MTT* is represented by the black dotted line (3 s) – anything above is an overestimation of the true *MTT*.

## Discussion

Dynamic susceptibility contrast (DSC) MRI is a powerful technique to assess perfusion in patients. However, ‘absolute’ perfusion values obtained with DSC have been suggested to not accurately reflect ground truth physiological parameters. For this reason, perfusion maps are typically investigated in diagnosis based on their relative distribution rather than their absolute values. In addition, it is not known whether the values obtained depend on the exact experimental conditions. To address these questions, we developed a novel signal framework for DSC MRI that incorporates signal contributions from intravascular and extravascular water proton spins at 3T for arterial, venous, and cerebral tissue voxels. This framework allowed us to model the MRI signal in response to changes in Gd and dOHb concentrations, and the effects that various experimental and tissue parameters have on perfusion quantification. We compared the simulated predictions to experimental data obtained at 3T on six healthy human subjects administered Gd and dOHb boluses in separate acquisitions. Based on our simulations and experimental findings, we found that perfusion quantification with Gd and dOHb depends on the parameters selected during acquisition and analysis. Our major findings in the simulations and experimental validations are as follows:

- Signal change in various cerebral regions (i.e., large vessels, GM, WM) scale inhomogeneously as a function of magnetic susceptibility values
- Increasing the simulated arterial *CBV* increases the estimated values of tissue *rCBV* and *rCBF*
- Reducing the baseline S_a_O_2_, increasing the susceptibility of the applied contrast agent (Gd *vs* dOHb), and/or increasing the ΔS_a_O_2_ reduces the estimated values of tissue *rCBV* and *rCBF*
- Normalizing tissue to venous rather than arterial signal increases the accuracy of tissue *rCBV* and *rCBF*
- Normalizing tissue to venous rather than arterial signal decreases the dependency of calculated tissue *rCBV* and *rCBF* on ΔS_a_O_2_, base-S_a_O_2_, and normalizing voxel *CBV*
- Shortening bolus duration increases the accuracy and reduces the estimated values of tissue *MTT*

### Signal Modeling Framework

The signal model applied in this work was previously used to simulate fMRI signal changes in response to neuronally-driven blood oxygenation changes and thus, changes in susceptibility (Uludag *et al*., 2009). However, given that the dOHb bolus used in this study is an externally driven susceptibility change of the same contrast agent as in fMRI, we could apply the same biophysical model to DSC MRI. To use this model for Gd contrast, we generalized the framework to Gd contrast as well. To do this, the relationship between Gd concentration and susceptibility changes had to be determined. In doing so, we found that at 3T, a 1 mM increase of Gd is equivalent in susceptibility to a 24.6% increase in dOHb (Eq. 8.2). Thus, Gd was modeled in the same way as dOHb, with differences only in the input concentration curve shape, duration, and susceptibility value. This concept is illustrated in Figures 1 and S3, which show that typical Gd and dOHb boluses occupy separate susceptibility ranges. That is, any difference in calculated physiological parameters (e.g., *CBV, CBF* and *MTT*) when using different contrast agents with a standard DSC model inversion can be majorly attributed to differences in the susceptibility and time course properties alone (of course, the differing compartmentalization of these contrast agents in vasculature yields small differences in quantification as well). For that reason, our model can be also applied to other contrast agents if their susceptibility value is known.

Our model calculates the tissue DSC MRI signal response to contrast agent resulting from both intravascular and extravascular relaxation rate changes in separate vascular compartments (artery, vein, capillary, arteriole, venule). This has theoretical advantages over previous *in vivo* and simulation work which models DSC MRI through a ‘bulk model’, wherein there is no theoretical separation between extravascular and intravascular compartments (Calamante *et al*., 2000; Chappell *et al*., 2015; Patil and Johnson, 2011). The effect of contrast agent on the proton spins in these unique compartments are not identical and should be treated as separate signal contributions (Uludag *et al*., 2009; Zhao *et al*., 2007; Blockley *et al*., 2008).

Kiselev *et al*. incorporates extravascular and intravascular signal contributions and notes differences in the calculated and ground truth *CBV, CBF*, and *MTT* of simulated voxels (Kiselev, 2001). By implementing a signal model accounting for both intravascular and extravascular contributions, Kiselev demonstrated that the contrast dosage can influence the calculated *CBV*, strengthening the case that a conventional DSC approach is not likely to yield absolute metrics, in agreement with our work (Kiselev, 2001). One of the main differences between our work and that performed by Kiselev is that the latter assumed that the intravascular relaxation rate is linearly dependent on the concentration of the contrast, in comparison to our model which assumes a quadratic relationship.

The intravascular transverse relaxation rate in response to contrast agent is somewhat contested. According to Lind *et al*., intravascular relaxation is linear with a relaxivity of 89 mM^-1^s^-1^ for Gd at 3T. Wilson *et al*. also reports a linear relationship and relaxivity of roughly 50 mM^-1^s^-1^ at 3T, dependent on the specific Gd-chelate. The linear relationships were ruled out in our work – for Lind *et al*., this value was determined *in vivo* and likely suffered high partial volume errors; the value 89 mM^-1^s^-1^ is far more reflective of extravascular relaxivity (Lind *et al*., 2020; Kjolby *et al*., 2006). For Wilson *et al*., the concentration of contrast agent was studied at levels beyond those typically observed (Wilson *et al*., 2017). On the other hand, numerous experimental and theoretical papers report quadratic dependencies between contrast agent (dOHb or Gd) and the intravascular relaxation rate (Akbudak *et al*., 2004; Blockley *et al*., 2008; Jensen and Chandra, 2000; Silvennoinen *et al*., 2003; van Osch *et al*., 2003; Zhao *et al*., 2007). As a linear intravascular relaxation rate would result in low or no dependency of perfusion values on the susceptibility value, our experimental results are a strong argument for a non-linear relationship of the contrast agent dose and intravascular relaxation rate (see below). Regarding the extravascular relaxation rate, there is better agreement based on rigorous Monte Carlo simulations that it is linearly related to the concentration of contrast agent (Boxerman *et al*., 1995; Kjolby *et al*., 2006; Uludag *et al*., 2009).

To the best of our knowledge, the two studies conducted by Kjolby *et al*. (Kjolby *et al*., 2006; Kjolby *et al*., 2009) are the only studies to simulate the transverse relaxation in tissue or artery from contrast agent in DSC MRI with a full signal model accounting for linear extravascular and quadratic intravascular relaxation rates. The paper in 2006 (Kjolby *et al*., 2006) focused on deriving tissue relaxivity in response to Gd and did not aim to quantify perfusion metrics, deviations from expected metrics, or quantification dependency on various tissue and contrast agent parameters. The paper in 2009 (Kjolby *et al*., 2009) set out to quantify perfusion metrics in response to Gd along with deviations from expected metrics when changing the partial volume level in the AIF. However, this work neglected separate vascular compartments in tissue and the extravascular signal magnitude contribution in the arterial voxel that arises from large vessel. Therefore, we believe that our model, incorporating both quadratic intravascular and linear extravascular components from separate vascular compartments, can serve as a more advanced model for DSC MRI simulations in future work.

### Signal Scaling Differs Between Gd and dOHb

While Gd-induced signal changes are larger than those induced by dOHb, signal scaling is not homogeneous in all areas of the brain (See Figures 6 and S4). When the dOHb signal change map is divided by the Gd signal change map, termed the *ΔS* ratio, we found a scaling difference between the large vessel (vein and artery) and tissue (GM and WM) voxels. Specifically, large vessel voxels show a larger *ΔS* ratio than tissue voxels in both simulations (Figure S4) and validations (Figure 3). The likely reason for this is due to non-linear signal scaling. When signal in the large vessels approaches zero, also known as saturation (Ellinger *et al*., 2000), signal change becomes non-linear with respect to dosage (doubling the dosage or *CBV* results in much smaller signal changes). Given that the signal from Gd contrast can approach the noise floor, not observed with a typical hypoxic bolus in our work or in the literature (Poublanc *et al*., 2021; Vu *et al*., 2021), there is an inherent difference between how signal is scaled between these two contrast agents in regions containing large vessels (more linear for dOHb and more non-linear for Gd, with respect to amount of contrast agent). This non-linear signal is taken into consideration when converting signal to relaxation rate, which accounts for the fact that relaxation rate and contrast agent concentration are within the exponential part of the signal equation. However, with the addition of noise in clinical data, conversion of signal to relaxation rate in the presence of signal saturation still results in inaccuracies – namely, overestimations of *CBV* and *CBF* (Ellinger *et al*., 2000).

Although of much smaller effect, there is also a scaling difference between the GM and WM voxels, wherein the GM *ΔS* ratio is lower than the WM *ΔS* ratio. This effect is observed in the validations (Figure 3) but not in the simulations (Figure S4). There are a few potential reasons why this may be the case. Firstly, in the validations, the mask designed for WM may incorporate more signal from draining vessel voxels that infiltrate the WM, in comparison to GM – in averaging the signal from WM and some vessels, the *ΔS* ratio would be increased. Another potential reason for the observed difference between GM and WM might be macrovascular contamination (Chappell *et al*., 2015), particularly the so-called blooming effect (reviewed by Willats and Calamante, 2012). It is known that Gd contrast results in a signal contamination in the cortical GM, where magnetic field disturbances from large vessels extends to the surrounding tissue (reviewed by Willats and Calamante, 2012). Assuming that these vessels are large arteries, this effect would be largely scaled down when using dOHb contrast due to the far lower signal arising from the arterial vessels on the cortical surface (refer to Figures 4 and 5). We did not simulate the blooming effect, which might be one reason that the simulations do not display a scaling difference between GM and WM. Further work would be required to isolate for and identify the level of the blooming effect when using Gd and dOHb. In summary, the signal change induced by Gd is larger than that induced by dOHb, but the scaling is not uniform throughout the brain, confirmed both in the experiments and simulations.

### Calculated rCBV and rCBF Depends on Simulated CBV of Normalizing Voxel, ΔS_a_O_2_, and base-S_a_O_2_

The simulations and *in vivo* tests reveal a dependence of *rCBV*, *rCBF*, and *MTT* on user-controlled parameters, such as the type of contrast agent, choice of normalizing voxel, ΔS_a_O_2_, base-S_a_O_2_, and bolus duration. Figure 5 shows that the area under the arterial bolus relaxation curve at 3T does not increase linearly as the simulated blood volume increases, but rather increases and then decreases at higher simulated *CBV* occupancy. This general pattern is also observed in previous work with Gd, where simulated arterial voxels with 80% *CBV* showed a greater signal change than those with 100% *CBV* (Kjolby *et al*., 2009). In simulated tissue voxels, the observed relaxation rate does not exhibit a maximum at intermediate CBV values for tissue (Figure 5). This results from the simulated tissue voxel being predominantly extravascular in composition. In contrast, it is the predominant intravascular component of simulated arterial and venous voxels that results in a non-linearity in *CBV* dependence (see Figure S1). As such, the model predicts that normalizing tissue voxels to an arterial voxel with high blood volume, counterintuitively results in higher calculated *rCBV* than when normalized to an arterial voxel with lower blood volume (Figure S1).

As shown in Figures 7 and 8, *rCBV* attained with dOHb becomes more accurate and decreases as ΔS_a_O_2_ increases and base-S_a_O_2_ decreases. This pattern is mainly observed when tissue is normalized to an arterial voxel with high *CBV*. It is the quadratic relaxation rate in the intravascular compartment alone that is responsible for these dependencies (Figure S1). The concept of the intravascular space contributing to overestimations is acknowledged in previous works (Kjolby *et al*., 2009; Wirestam *et al*., 2010). While it will be important to control for ΔS_a_O_2_ and base-S_a_O_2_ across scans to maintain precision, this only applies when tissue is normalized to an arterial voxel with a large amount of intravascular space (roughly 80-100%), as evident in Figure S1.

Bleeker *et al*. (Bleeker *et al*., 2009) showed that it is beneficial to select an AIF adjacent to the intravascular space – our work supports this claim for two reasons. Firstly, our simulations show that normalizing tissue to an AIF with a lower simulated *CBV* (Figure S1) results in higher accuracy of the calculated *rCBV*. Secondly, our simulations reveal a lower dependence of the calculated *rCBV* on ΔS_a_O_2_ and base-S_a_O_2_ (Figure S1) when tissue is normalized to an AIF with a lower simulated *CBV*.

Wirestam *et al*. (Wirestam *et al*., 2010) has shown that doubling [Gd] at 3T, increasing the susceptibility of contrast agent, increases the calculated *rCBV* and *rCBF* – they suggest that this counterintuitive finding is likely due to partial volume errors and signal saturation. We show that increasing the susceptibility of contrast agent decreases the calculated *rCBV* and *rCBF*, without the confound of partial volume errors in the case of our simulations. Of note, given that *rCBV* acquired with Gd (gadodiamide) in a GRE scan overestimates the true *CBV* by roughly three times (Østergaard *et al*., 1998; Simonsen *et al*., 2000), *rCBV* obtained with Gd does not represent a ground truth. Therefore, normalizing our dOHb *rCBV* results to Gd *rCBV* results (Figure 8) does not imply normalizing to the true *CBV*.

### Calculated rCBV and rCBF Depends on the Choice of Artery or Vein for Tissue Normalization

From a clinical perspective, it might be beneficial to normalize tissue signal to a venous as opposed to arterial voxel, as the larger radius of the venous blood vessel results in less partial volume errors (reviewed by Willats and Calamante, 2012). Although there are still limitations to this type of analysis in DSC MRI (Knutsson *et al*., 2007), we sought to investigate how perfusion quantification depends on arterial *vs* venous normalization.

In simulations and *in vivo* tests, normalizing 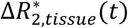 to 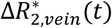 results in lower overestimations of *rCBV* and *rCBF*, along with a reduced ΔS_a_O_2_ and base-S_a_O_2_ dependency in comparison to when 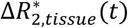 is normalized to 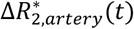 (Figures 7, 8, and S1). As previously described, a lower base-S_a_O_2_ results in larger relaxation rate changes on account of the quadratic intravascular dependence on SO_2_, which then influences perfusion estimation. Thus, if a vein and an artery are provided the same ΔS_a_O_2_ and are simulated with the same *CBV*, it is clear from the previous results that the voxel with a lower base-S_a_O_2_, as is the case in veins, will have a larger relaxation rate change (Figure 4). This theoretical discovery likely contributes to the high venous to arterial relaxation rate peak ratio observed in previous studies using dOHb in DSC MRI (Poublanc *et al*., 2021; Vu *et al*., 2021). When tissue signal is then normalized to the venous voxel with a larger relaxation rate change relative to artery, and thus, larger area under curve, a lower and more accurate estimate of *rCBV* and *rCBF* is obtained (Refer to Eq. 10 and Figure 8). At this higher baseline susceptibility in veins, due to an already low base-S_a_O_2_, we found that varying the ΔS_a_O_2_ and base-S_a_O_2_ through different paradigms has less of an effect on perfusion quantification in both the simulations and experimental validations. A reduced contrast dosage dependency and lower calculated *rCBV* when normalizing to vein as opposed to artery is observed in previous experimental results as well (Wirestam *et al*., 2010).

Our experimental findings that show a link between contrast agent parameters and perfusion quantification supports a quadratic, as opposed to linear, intravascular relaxation rate in response to contrast agent. If this relationship were linear, then modifying base-S_a_O_2_ from one scan to the next would not influence perfusion quantification, as the base-S_a_O_2_, for example, would cancel out with its effect at baseline, as is observed in the extravascular compartment (Refer to Eqs. 4–5). In other words, the fact that the *rCBV* and *rCBF* values obtained from the experimental data depend on various experimental and analysis choices is a strong indication that the intravascular relaxation rate is not linearly related to the concentration of the contrast agent. This implies that a standard DSC MRI analysis will not yield absolute values of perfusion quantification, whereas a model that accounts for these relationships, such as the one employed in our work, should provide more accurate estimations.

### Calculated MTT and rCBF Depends on Bolus Duration

The experimental data reveal an overestimation of the calculated *MTT* in all participants when using dOHb in comparison to Gd for quantification (Figure 9). Vu *et al*. (Vu *et al*., 2021) also reported high calculated *MTT* values when using dOHb as a contrast agent in a standard DSC MRI analysis. We consulted the simulations and literature to determine the differences between the properties of Gd and dOHb boluses that contribute to this discrepancy.

The simulations reveal that bolus duration influences *MTT* calculation, which in turn affects *rCBF* (Figure 9). This is expected as *MTT* becomes small relative to the duration and, hence, the amount of data encoding *MTT* occupies less of the overall bolus duration. We also found that the thresholding of singular values causes a duration dependency – longer boluses resulted in the removal of more singular values from the AIF. It is known that as more singular values are removed, there is a greater underestimation of *rCBF* and greater overestimation of *MTT* (Bjornerud and Emblem, 2010). Thus, a Gd bolus, with shorter duration and less removed singular values, results in the calculation of a lower and more accurate *MTT* relative to the *MTT* calculated from a dOHb bolus. In our simulation work, when the dOHb bolus is shortened to the duration of Gd, the calculated *MTT* is more similar to that obtained with Gd (Figure 9).

Although noise was not directly simulated in our work, it is known that *SNR* is another major parameter influencing *MTT* calculation, wherein more singular values from the AIF matrix are removed when there is a lower *SNR*, resulting in a greater underestimation of *rCBF* and overestimation of *MTT* (Willats and Calamante, 2012). Thus, a combination of low *SNR* and long bolus duration results in higher estimations of *MTT* than the combination of high *SNR* and short bolus duration. Given that hypoxic boluses are of longer duration and lower *SNR* than Gd boluses in our experiments, it is especially the case that we might expect to observe a lower calculated *MTT* when using the Gd bolus.

Previous work where Gd was doubled in concentration revealed an increase in calculated *MTT* relative to a standard dosage (Wirestam *et al*., 2010) – assuming the bolus had increased in duration due to the bolus doubling, our findings help explain this observation.

### Considerations for Clinical Implementation of dOHb and Gd

For dOHb contrast to attain an *SNR* like that of Gd in a single bolus, the oxygen saturation must drop below physiological tolerance (Figure 1 and Table S1). However, DSC using dOHb has several advantages (Poublanc *et al*., 2021; Vu *et al*., 2021). Firstly, as dOHb is an endogenous contrast agent, it does not share the safety limitations associated with Gd. As well, [dOHb] does not saturate, recirculate, or accumulate, as the blood oxygenation is reset to a pre-determined value in the lungs using the RespirAct. In contrast to Gd, the concentration of dOHb in the arterial blood can be determined from end-tidal gas analysis which may be useful as a precise AIF and aid in quantitative modeling (Fierstra *et al*., 2011; Ito *et al*., 2008). Finally, dOHb acquisitions can be repeated and averaged to improve *SNR* closer to that of Gd (‘Average dOHb’ in Table S1).

While the smaller hypoxic challenge (a drop in S_a_O_2_ from 100% to 90%) administered in our study is equivalent to experiencing reduced S_a_O_2_ in high altitudes (e.g., in Aspen, Colorado), the larger drop (100% to 75% S_a_O_2_) is closer to the S_a_O_2_ in the Everest Base Camp (Rojas-Camayo *et al*., 2018). Individuals may spend many months in these environments without additional oxygen or altitude sickness, minimizing the safety concern in our study where hypoxia only lasts for roughly 30 s.

### Limitations and Directions

One notable limitation in the signal modeling was the uncertainty of the maximum concentration of Gd for input into the simulations. Researchers have reported a vast array of values for this (Kellner *et al*, 2018; Lind *et al*, 2020; Patil *et al*., 2013); therefore, we sought to balance two competing concerns: creating a concentration profile for Gd which results in tissue 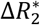 curves representative of those in our validation data; avoiding signal saturation, or hitting the noise floor, in arterial and venous voxels. A further limitation arises with respect to potential vasodilation from the hypoxic stimulus. Previous work determined that dropping the S_a_O_2_ from 100% to 90% for a 20-minute duration is the threshold for statistically significant cerebral vasodilation in humans (Gupta *et al*., 1997). According to more recent work, an S_a_O_2_ of approximately 70% results in increases to the *CBF* (Mardimae *et al*., 2012). However, the vascular response is delayed for about 3 minutes following hypoxia (Harris *et al*., 2013). Thus, as we provided transient drops in S_a_O_2_ to 75%, at the lowest, for approximately 30 s, we expect that significant vasodilation is avoided in our work. Nevertheless, our model can also account for vasodilation if *CBV* changes are known or can be estimated.

Future work with dOHb contrast should consider the use of a higher field strength scanner to further bolster *SNR* – this might be the best way to maximize the benefits of dOHb as long as the arterial signal does not become saturated. The use of 7T, for example, might circumvent the quadratic intravascular contribution, as it is known that at higher field strengths, the intravascular contribution in T2*-weighted acquisitions is heavily reduced relative to extravascular signal contribution (Uludag *et al*., 2009). Another potential direction would be to investigate the use of dOHb contrast in clinical conditions such as brain tumors, steno-occlusive vascular disease, and dementia. This could first be simulated with our framework and then validated with clinical scans. In addition, it would be useful to explore a model-based DSC MRI analysis method by inputting the subject’s S_a_O_2_ profile, as opposed to a manually selected AIF, into the signal framework described in our work – this might allow for more consistent quantification as there is no need to threshold singular values or select an AIF (Poublanc *et al*., 2021).

## Conclusion

In this work, we implemented a signal framework to study how various contrast agent parameters and tissue properties influence perfusion estimation and applied this model to simulate DSC MRI signal in response to Gd and dOHb contrast agents. In doing so, perfusion quantification dependencies were revealed that agreed in the simulations and *in vivo* experiments. We hope that this model can inform future research and clinical protocols that employ dOHb and Gd as contrast agents.

## Acknowledgments

We would like to thank Dr. Jean J. Chen for valuable feedback on a previous version of the manuscript. The research conducted in this paper was supported by funding from the Canadian Institutes of Health Research (CIHR). The study was also supported by the Institute for Basic Science, Suwon, Republic of Korea (IBS-R015-D1) to Kamil Uludag.

## Supplementary Figure Captions

**Table S1. Summary of Perfusion Data Collected with dOHb and Gd Contrast Agents.** Average and standard deviations are pooled from a total of six healthy participants (age 22-60; one female). Low baseline dOHb is only pooled from five participants due to motion.

**Figure S1. dOHb *rCBV* as a Function of Simulated Arterial or Venous *CBV* (%).** *rCBV* accuracy (y-axis) for tissue (*CBV* = 4%) when normalized to artery (or vein) of varying simulated *CBV* (x-axis). *rCBV* accuracy calculated as *rCBV_calculated_* divided by *rCBV_simulated_*. All values above 1 (dotted line) overestimate the *rCBV_simulated_*. **Left.** Three scenarios are described in this figure: only extravascular component of signal model used in simulated artery (black plus); only intravascular component of signal model used in simulated artery (dark grey square); full signal model used in simulated artery (light gray diamond). **Right.** Tissue data normalized to AIF (black) or VOF (grey). Four scenarios are described in this figure: large ΔS_a_O_2_ with high base-S_a_O_2_ (diamond), medium ΔS_a_O_2_ with high base-S_a_O_2_ (square), small ΔS_a_O_2_ with high base-S_a_O_2_ (plus), small ΔS_a_O_2_ with low base-S_a_O_2_ (triangle).

**Figure S2. GM, WM, and Large Vessel Masks Created in Functional Space. Red.** Large vessels. **Blue.** GM. **White.** WM. For WM, voxels containing values within the lowest 15-20% of the ΔS data range were kept; for GM, the highest 75% were kept. The maps were binarized and linearly registered to the functional space of each dOHb paradigm. GM and WM were further masked with a thresholded *ε_t_* map (voxels with values in the lowest 10% of data range were kept in mask) to remove noisy voxels in peripheral brain tissue, skull base, vein, and artery. The GM and WM regions were finally masked with a thresholded Gd *MTT* map (lowest 70% of robust range voxels were kept in mask) to remove voxels in the CSF. Large vessel mask (only used for results in Figure 5) generated by taking the highest 50% of *ΔS_Gd_* or *ΔS_dOHb_* robust range.

**Figure S3. Perfusion Ratios as a Function of Peak Susceptibility Induced by Contrast Agent.** The region shaded in blue is the typical peak susceptibility elicited in a hypoxic dOHb bolus (Peak ΔS_a_O_2_ ≈ 10-25%). The region shaded in red is the typical peak susceptibility elicited in a Gd bolus (Peak [Gd] ≈ 2-5 mM). **Left.** rCBV (%) as a function of peak susceptibility induced by contrast agent. Simulated artery *CBV* = 100%; simulated tissue *CBV* = 4%. **Right.** Ratio of arterial to venous relaxation rate time integral as a function of peak susceptibility induced by contrast agent. Simulated arterial and venous *CBV* = 100%.

**Figure S4. ΔS_dOHb_/ΔS_Gd_ for Simulated Arterial (red), Venous (blue), GM (grey), and WM (white) Voxels.** Simulated dOHb bolus followed a 98% to 84% S_a_O_2_ profile. Simulated Gd bolus attained a peak concentration of 6 mM. Simulated *CBVs* were as follows: artery (90%), vein (90%), GM (4%), and WM (2%).

**Table S2. Table of Parameters**

