## Supplementary Data for "Perfusion Quantification in the Human Brain Using DSC MRI – Simulations and Validations at 3T"

### Slide 1
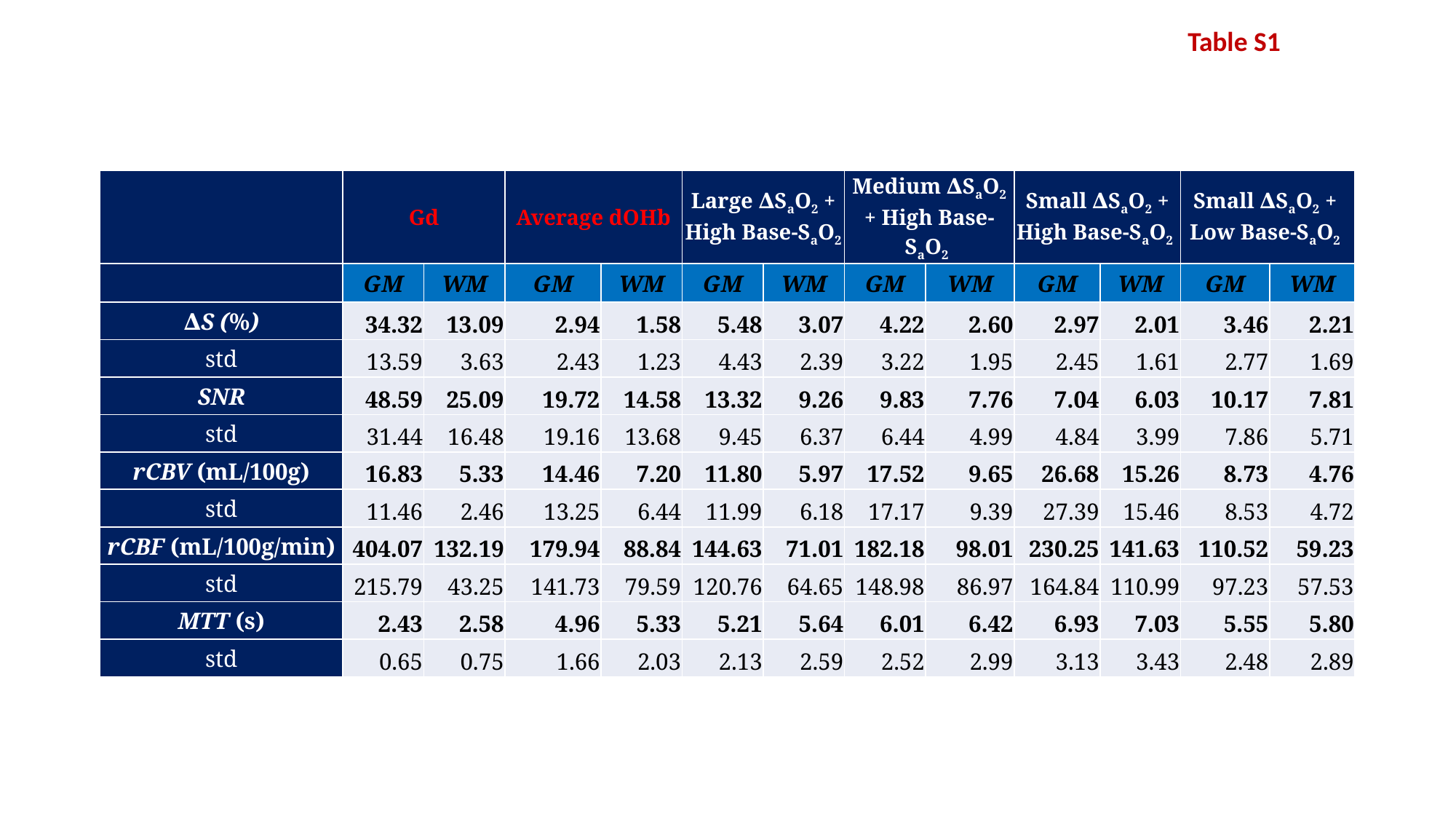

Table S1
| | Gd | | Average dOHb | | Large ∆S_a_O_2_ + High Base-S_a_O_2_ | | Medium ∆S_a_O_2_ + High Base-S_a_O_2_ | | Small ∆S_a_O_2_ + High Base-S_a_O_2_ | | Small ∆S_a_O_2_ + Low Base-S_a_O_2_ | |
| --- | --- | --- | --- | --- | --- | --- | --- | --- | --- | --- | --- | --- |
| | GM | WM | GM | WM | GM | WM | GM | WM | GM | WM | GM | WM |
| ∆S (%) | 34.32 | 13.09 | 2.94 | 1.58 | 5.48 | 3.07 | 4.22 | 2.60 | 2.97 | 2.01 | 3.46 | 2.21 |
| std | 13.59 | 3.63 | 2.43 | 1.23 | 4.43 | 2.39 | 3.22 | 1.95 | 2.45 | 1.61 | 2.77 | 1.69 |
| SNR | 48.59 | 25.09 | 19.72 | 14.58 | 13.32 | 9.26 | 9.83 | 7.76 | 7.04 | 6.03 | 10.17 | 7.81 |
| std | 31.44 | 16.48 | 19.16 | 13.68 | 9.45 | 6.37 | 6.44 | 4.99 | 4.84 | 3.99 | 7.86 | 5.71 |
| rCBV (mL/100g) | 16.83 | 5.33 | 14.46 | 7.20 | 11.80 | 5.97 | 17.52 | 9.65 | 26.68 | 15.26 | 8.73 | 4.76 |
| std | 11.46 | 2.46 | 13.25 | 6.44 | 11.99 | 6.18 | 17.17 | 9.39 | 27.39 | 15.46 | 8.53 | 4.72 |
| rCBF (mL/100g/min) | 404.07 | 132.19 | 179.94 | 88.84 | 144.63 | 71.01 | 182.18 | 98.01 | 230.25 | 141.63 | 110.52 | 59.23 |
| std | 215.79 | 43.25 | 141.73 | 79.59 | 120.76 | 64.65 | 148.98 | 86.97 | 164.84 | 110.99 | 97.23 | 57.53 |
| MTT (s) | 2.43 | 2.58 | 4.96 | 5.33 | 5.21 | 5.64 | 6.01 | 6.42 | 6.93 | 7.03 | 5.55 | 5.80 |
| std | 0.65 | 0.75 | 1.66 | 2.03 | 2.13 | 2.59 | 2.52 | 2.99 | 3.13 | 3.43 | 2.48 | 2.89 |

### Slide 2
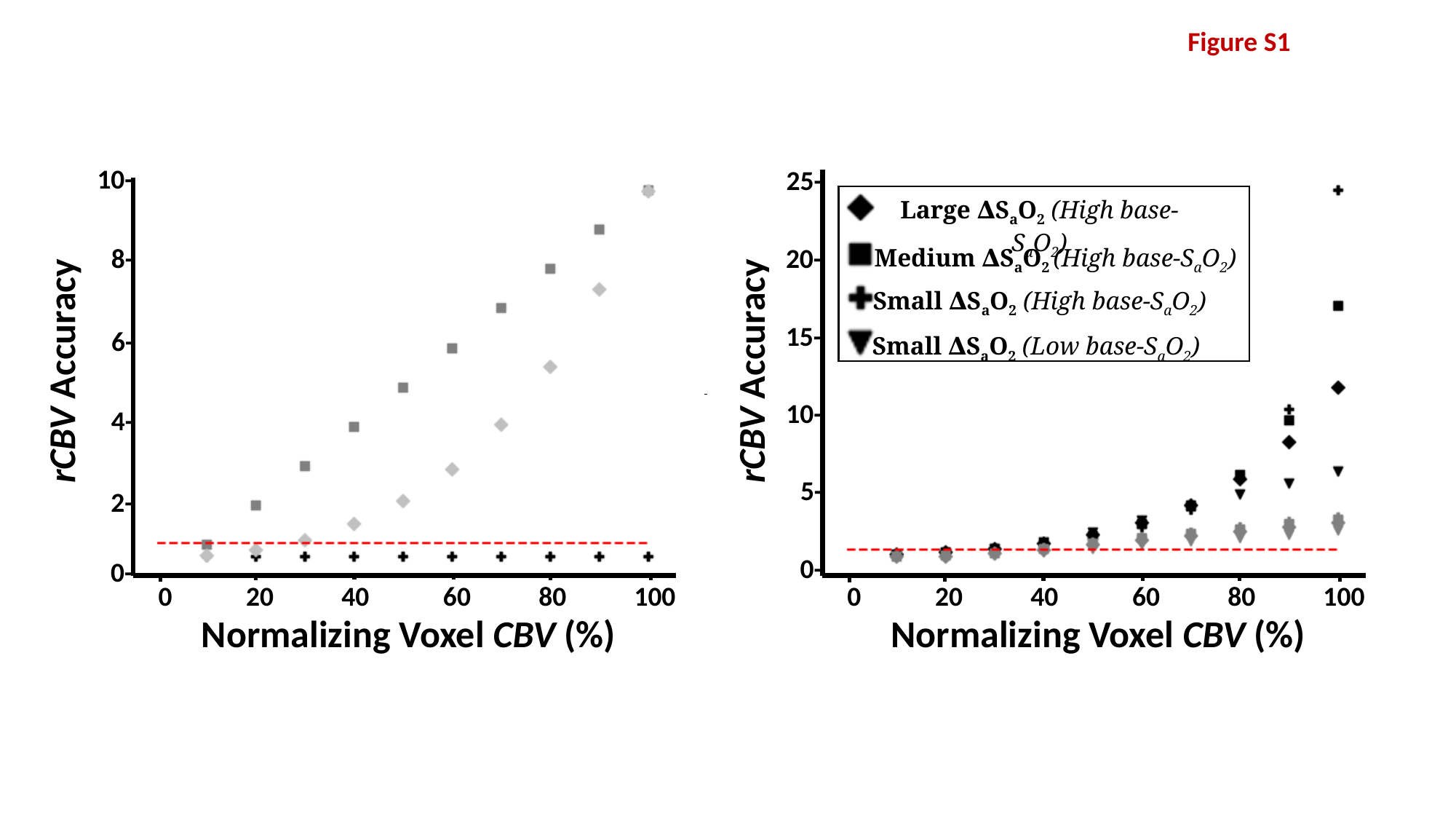

Figure S1
10-
25-
Large ∆S_a_O_2_ (High base-S_a_O_2_)
20-
8-
Medium ∆S_a_O_2_ (High base-S_a_O_2_)
Small ∆S_a_O_2_ (High base-S_a_O_2_)
15-
6-
Small ∆S_a_O_2_ (Low base-S_a_O_2_)
rCBV Accuracy
rCBV Accuracy
10-
4-
5-
2-
0-
0-
0 20 40 60 80 100
0 20 40 60 80 100
Normalizing Voxel CBV (%)
Normalizing Voxel CBV (%)

### Slide 3
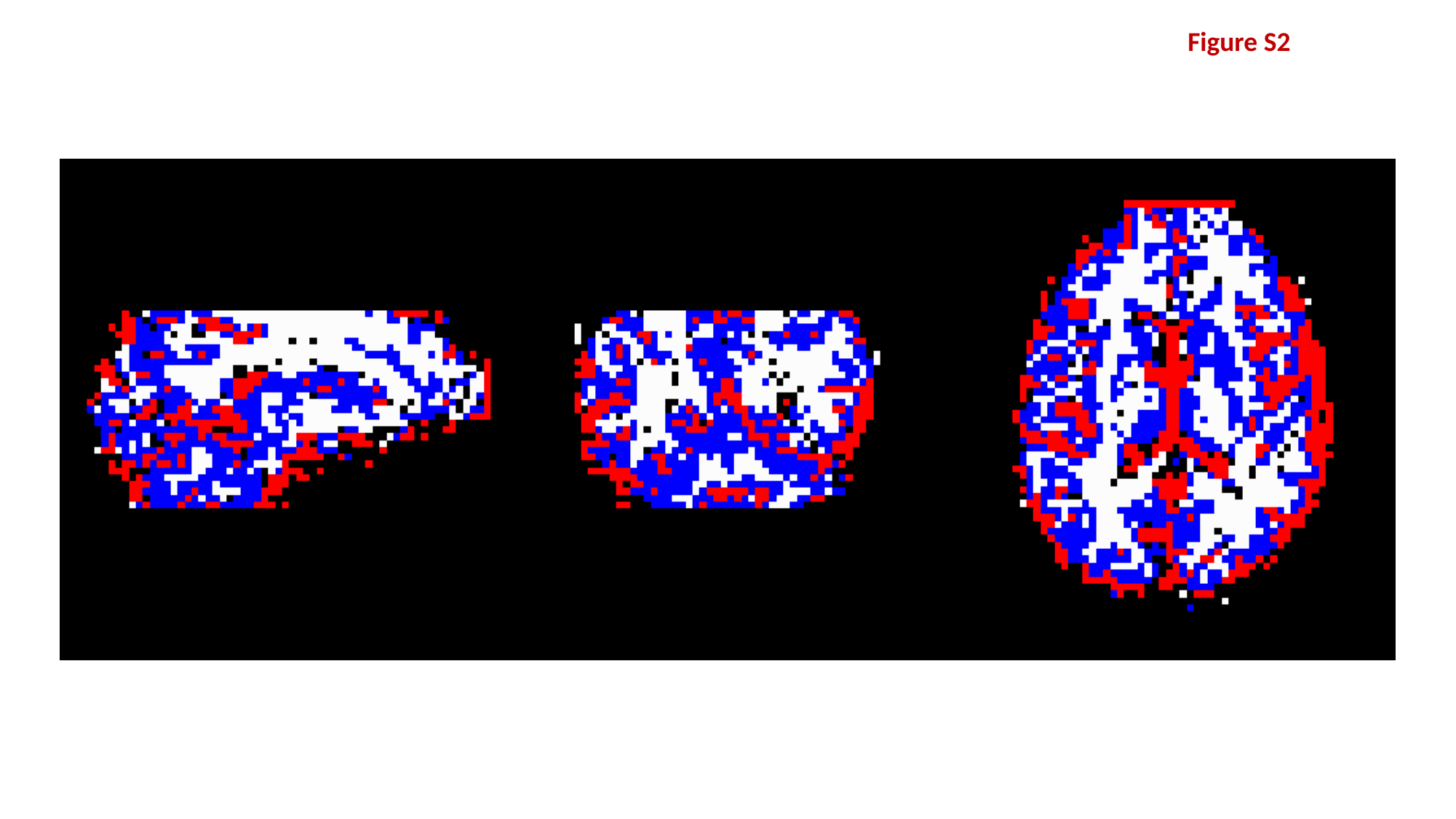

Figure S2

### Slide 4
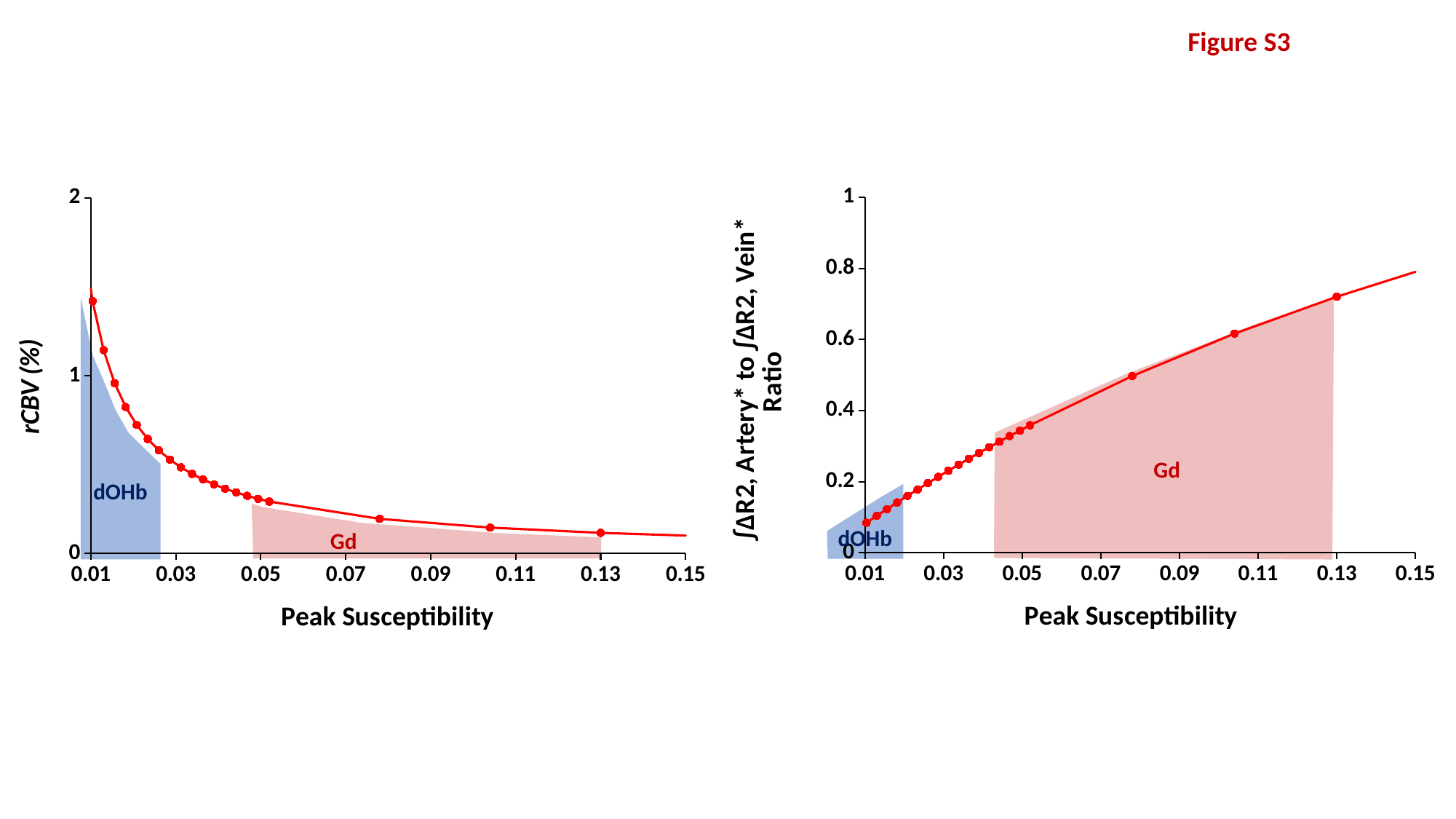

Figure S3
#### Chart
| Category | ∫Arterial to ∫Venous Ratio |
|---|---|
#### Chart
| Category | CBV |
|---|---|
dOHb
Gd
dOHb
Gd

### Slide 5
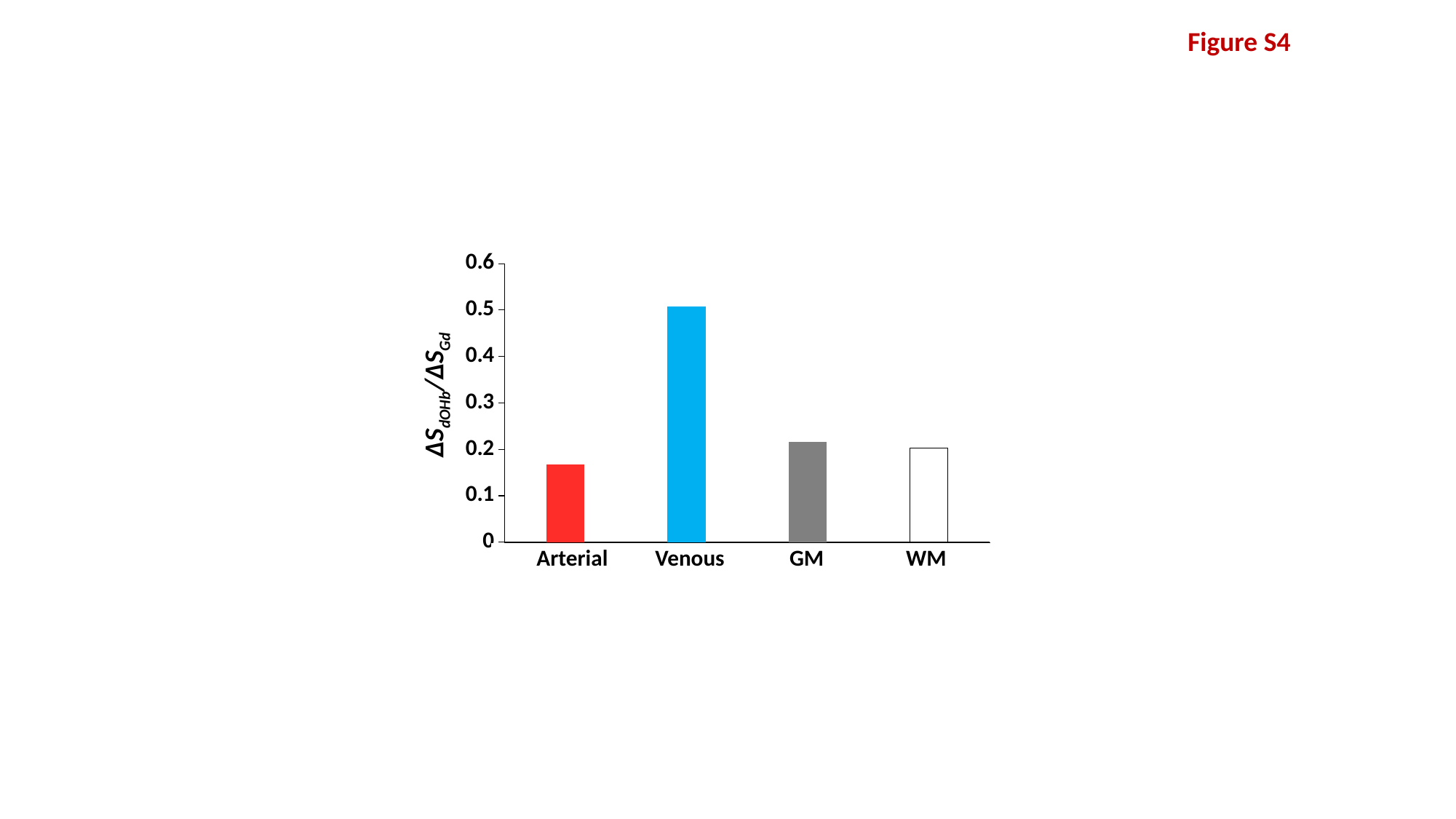

Figure S4
#### Chart
| Category | ∆Sdohb/∆Sgd |
|---|---|
| Artery (CBV = 90%)) | 0.16820000000000002 |
| Vein (CBV = 90%)) | 0.5075 |
| GM (CBV = 4%) | 0.2162892869212572 |
| WM (CBV = 2%) | 0.20299812617114302 |∆S_dOHb_/∆S_Gd_
WM
GM
Arterial
Venous

### Slide 6
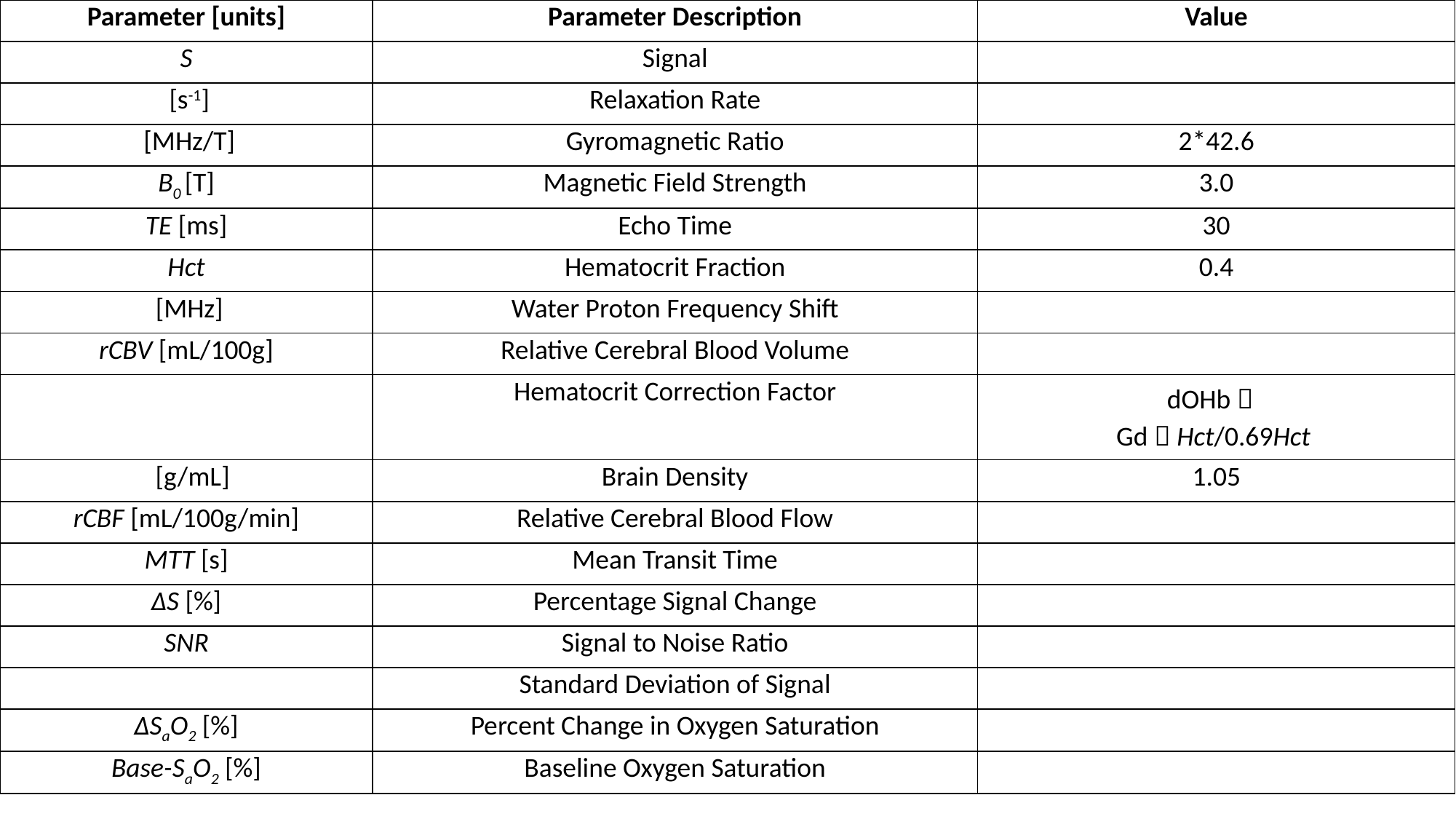

Table S2
